# Sex-dependent interferon signaling drives female-biased vulnerability in Alzheimer’s disease

**DOI:** 10.1101/2025.08.22.671724

**Authors:** Verónica López-López, Gerard Iniesta, Marcos Galán-Ganga, Alejandro Expósito-Coca, Aysha Bhojwani-Cabrera, Marina Guillot-Fernández, Albert Giralt, José P. López-Atalaya

**Author notes:** Correspondence: José P. López-Atalaya, Instituto de Neurociencias, Consejo Superior de Investigaciones Científicas, Universidad Miguel Hernández, Alicante, Spain.

## Abstract

Alzheimer’s disease (AD) disproportionately affects women, yet the neurobiological mechanisms underlying this sex bias remain poorly understood. Here, we identify a sex-dependent activation of type I interferon (IFN-I) signaling as a contributor to this disparity. Transcriptomic profiling of brain from AD patients revealed selective enrichment of IFN-I pathway in females. This immune signature was mirrored in the APP/PS1 mouse model of AD, where females exhibited more pronounced amyloid accumulation, neuroinflammation, and neurodegeneration. Acute IFN-I activation reproduced pathological features of AD, whereas chronic IFN-I elevation in APP/PS1 mice aggravated disease progression. In contrast, pharmacological targeting of IFN-I response by inhibiting the cGAS-STING pathway in APP/PS1 female mice reduced neuropathological burden, and preserved cognitive performance. Together, these findings identify interferon signaling as a modifiable and sex-linked driver of AD pathology. Our study uncovers a critical neuroimmune mechanism contributing to female-biased vulnerability and highlights interferon modulation as a promising therapeutic strategy in AD.

## Introduction

Alzheimer’s disease (AD) is a progressive and ultimately fatal neurodegenerative disorder and the most common cause of dementia worldwide (1–3). With an aging global population, the number of individuals affected by AD is projected to exceed 100 million by 2050, placing an enormous burden on healthcare systems and caregivers (1, 3–5). The neuropathological hallmarks of AD - extracellular amyloid-β plaques, intracellular tau neurofibrillary tangles, and chronic neuroinflammation - are accompanied by cognitive and behavioral impairments that progressively erode autonomy and quality of life (2, 3, 6). Despite decades of intense research, currently approved treatments provide only modest symptomatic relief, and no curative or disease-modifying therapies have yet been established (6–8).

A consistently reported yet underexplored feature of AD is its disproportionate impact on females. Females comprise nearly two-thirds of individuals diagnosed with AD (1, 3, 5), and lifetime risk estimates indicate that approximately 1 in 5 females, compared to 1 in 10 males, will develop the disease (1, 4). Although increased longevity contributes to this imbalance, it does not fully account for the observed differences (9–12). Clinical and neuroimaging studies have revealed that females with AD often exhibit faster cognitive decline, greater neuropathological burden, and more rapid progression from mild cognitive impairment to dementia than their male counterparts (9, 13–18). Biological sex thus acts as an independent modifier of AD risk and trajectory, shaping both susceptibility and disease course.

Despite its clinical significance, the mechanistic basis for sex-related differences in AD remains poorly defined. Historically, biological sex has been underrepresented or insufficiently analyzed in both clinical trials and experimental models (19–22). Many preclinical studies rely exclusively on male animals, or include both sexes without stratified analysis, thereby missing potentially critical sex-dependent effects (20, 23–25). This limitation not only undermines the translational relevance of preclinical findings but also hinders the development of sex-informed therapeutic strategies.

Addressing this knowledge gap requires a targeted focus on molecular and cellular pathways that are both central to AD pathogenesis and modulated by sex. In this regard, the innate immune system - particularly neuroimmune signaling within glial networks - has emerged as a key candidate. Genome-wide association studies (GWAS) have identified numerous AD susceptibility loci involved in immune regulation, including variants in *TREM2*, *CD33*, and *HLA* genes, among others (26–29). These discoveries have expanded the conceptual framework of AD beyond classical amyloid and tau cascades, implicating immune dysregulation and glial biology as core features of disease pathogenesis (30–34).

In parallel, a growing body of research has shown that immune responses are strongly shaped by biological sex. Females generally exhibit stronger baseline and stimulus-induced immune responses, both peripherally and in the central nervous system (35–39). While this amplified responsiveness may enhance pathogen clearance, it is also linked to greater vulnerability to chronic inflammation and autoimmunity (35, 36, 38). In the context of neurodegeneration, sex-biased immune traits may alter the threshold, magnitude, or persistence of neuroinflammatory responses, potentially influencing susceptibility to and progression of AD (14, 40–42).

Whether such immunological differences translate into distinct molecular trajectories or pathological outcomes in AD remains an open question. Biological sex may shape diverse cellular programs - including glial activation states (43–46), neuronal vulnerability, and synaptic remodeling (47–50) - that collectively influence disease dynamics. However, these hypotheses require systematic and mechanistic investigation. Understanding how sex interacts with disease-relevant pathways is essential for explaining observed clinical disparities and for identifying novel therapeutic targets that reflect this dimension of disease biology.

In this study, we sought to investigate the molecular mechanisms underlying sex-associated differences in AD vulnerability. We combined transcriptomic analysis of *postmortem* human brain tissue with functional studies in established mouse models of AD. By stratifying human samples by sex, age, and brain region, we identified a pronounced activation of type I interferon (IFN-I) signaling in females. To explore the causal role of sex-dependent IFN-I signaling in disease pathogenesis, we employed both pharmacological and genetic manipulations *in vivo* and assessed the impact of modulating IFN-I signaling on neuroinflammation, neuropathology, and cognitive performance. This integrative approach establishes IFN-I signaling as a key contributor to sex-associated differences in AD vulnerability and highlights its potential as a therapeutic target.

## Results

### Sex Differences in Interferon Pathway Activation Are Associated with Increased Alzheimer’s Risk and Prevalence in Women

Epidemiological data from the Alzheimer’s Facts and Figures 2024 report highlight a markedly increased lifetime risk of AD in women (51). At both 45 and 65 years of age, the estimated lifetime risk of developing AD is approximately 20% for women versus 10% for men (51–54). Among individuals aged 65 and older, women account for approximately 61% of all AD cases, while men account for 39% (51, 53, 55). These sex disparities in disease prevalence and risk are illustrated schematically (Figure 1A).

**Figure 1.**
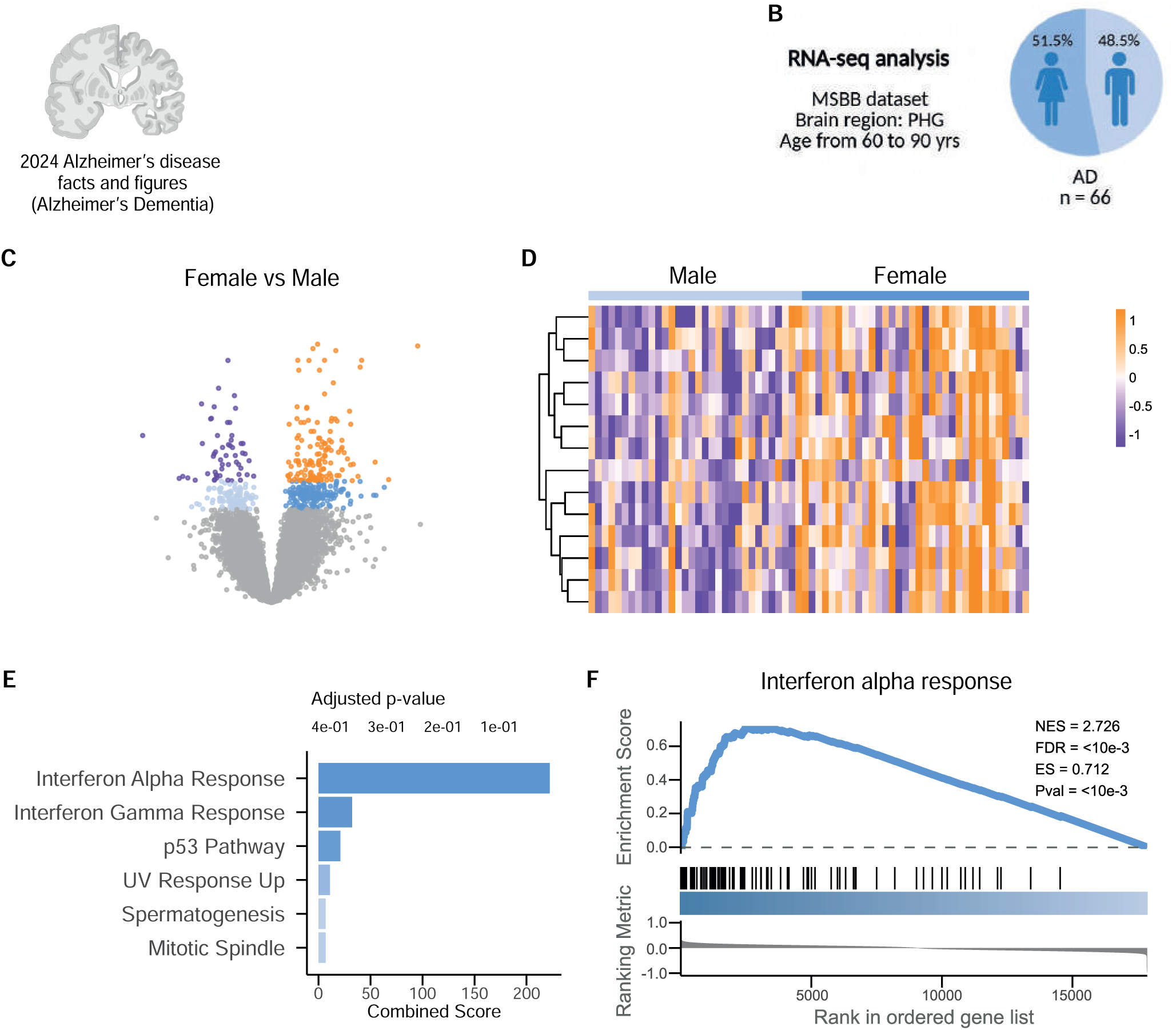
Enhanced interferon signaling associates with female-biased Alzheimer’s disease risk and prevalence. (A) Diagram illustrating lifetime risk and prevalence of AD, with women showing higher risk (20% vs. 10%) and prevalence (61% vs. 39%) compared to men (Alzheimer’s Facts and Figures 2024). (B) Cohort overview of RNA-sequencing data from the MSBB, comprising 34 female and 32 male AD patients (aged 60–90 years) for parahippocampal gyrus analysis. (C) Volcano plot of differential gene expression in female vs. male AD brains. Significantly upregulated genes (FDR < 0.05) are shown in orange, and genes with 0.05 ≤ FDR < 0.1 are shown in dark blue. Interferon alpha signaling genes are highlighted. (D) Heatmap showing elevated expression of the top differentially expressed (FDR < 0.1) interferon-related genes in female AD samples compared to male samples. (E) Functional enrichment analysis of significantly upregulated genes (FDR < 0.05) in female AD brains showing enrichment of interferon-alpha and interferon-gamma response pathways. (F) GSEA plot showing coordinated upregulation of the interferon alpha response pathway in female AD brains.

While several clinical studies have reported faster disease progression and greater pathological burden in females (13, 14, 16, 17), the molecular mechanisms driving this increased vulnerability remain poorly understood. To address this, we leveraged RNA-sequencing data from the parahippocampal gyrus (PHG) of AD patients in The Mount Sinai Brain Bank (MSBB) study (56). Since women predominate at the most advances ages (>90 years) due to their longer survival, we restricted our analysis to individuals aged 60 to 90 years to minimize potential confounding, yielding a final cohort of 34 females and 32 males AD patients (Figure 1B, Supplemental Figure 1, A and B, and Supplemental Table 1). Initial analysis of these samples still revealed an age effect across sexes (Supplemental Figure 1, C-F). To account for this potential confounder, age was included as a covariate in all subsequent analyses.

Differential gene expression analysis revealed a prominent female-biased transcriptional signature related to interferon signaling in AD patients, including genes such as *GBP4, CNP, PSMB9, PSMB8, EPSTI1, IFI44L, and TRIM26* (Figure 1, C and D, and Supplemental Table 1). Functional enrichment analysis revealed that “Interferon Alpha Response” was the most significantly enriched “hallmark” gene set (57) in females, followed by “Interferon Gamma Response” while “p53 Pathway” showed borderline enrichment (adj. p = 0.05) (Figure 1E). Gene Set Enrichment Analysis (GSEA) (58) further confirmed strong and coordinated upregulation of the interferon alpha response pathway in female AD samples (Figure 1F).

To place these sex-stratified findings in the broader context of AD pathology, we compared AD versus age-matched control samples without stratifying by sex (n = 108; 42 controls and 66 AD) (Supplemental Table 1). In this pooled analysis, functional profiling identified “Epithelial Mesenchymal Transition” as the top enriched pathway, followed by “Interferon Gamma Response” and “Interferon Alpha Response” (Supplemental Figure 1G), in addition to other classical AD-associated pathways such as ‘Cholesterol Homeostasis’ (Supplemental Table 1). However, stratifying by sex revealed a marked shift in the dominant transcriptional programs: in males (n = 55; 23 controls and 32 AD), “Epithelial Mesenchymal Transition” emerged as the principal upregulated pathway, whereas in females (n = 53; 19 controls and 34 AD), “Interferon Alpha Response” became the leading enriched pathway with “Epithelial Mesenchymal Transition” ranking lower (Supplemental Figure 1, H and I). This divergence underscores the critical influence of sex on the molecular architecture of AD, revealing that the primary immune program engaged by the disease differs fundamentally between males and females.

Next, we examined whether this transcriptional signature was also detectable under non-pathological conditions. To this end, we analyzed transcriptomic data obtained from the PHG of age-matched control individuals (n = 42; 19 females and 23 males) (Supplemental Table 1). Functional analysis of sex-associated differentially expressed genes revealed no significant enrichment of interferon-related pathways (Supplemental Figure 1, J and K). GSEA did not reveal significant enrichment of interferon-related pathways (Supplemental Figure 1L), indicating that baseline differences between males and females are minimal, and that the presence of AD pathology is required to unmask the robust sex-dependent interferon signature.

Together, these findings show that female AD patients exhibit a sustained upregulation of IFN-I in the parahippocampal gyrus. This persistent immune activation may contribute to the greater vulnerability and higher prevalence of AD observed in women, highlighting sex-specific neuroimmune mechanisms as potential contributors to disease susceptibility and progression.

### Sex Differences in Amyloid Pathology and Neurodegeneration in APP/PS1 Mice

To assess whether the sex differences observed in human AD are recapitulated in mouse models, we analyzed male and female APPswe/PSEN1dE9 (APP/PS1) transgenic mice. This model develops robust Aβ pathology and neuroinflammatory changes. Histological analyses were performed at 6 months of age, a stage when amyloid deposition is actively progressing in the hippocampus and cortex of APP/PS1 mice (Figure 2A). Immunohistochemical staining with the anti–amyloid-β (Aβ) antibody 6E10 revealed significantly higher plaque burden in both the cortex and hippocampus of female mice compared with males (Figure 2B). Thioflavin T staining confirmed these observations, showing more extensive fibrillar amyloid deposits in females (Figure 2C). Consistently, immunostaining with the OC antibody, which recognizes β-sheet-rich Aβ conformations, revealed greater signal in female brains (Figure 2D), further validating increased pathological aggregation. Notably, plaque size was similar between sexes (Supplemental Figure 2, A-C), suggesting that the sex difference primarily reflects increased plaque number or density rather than altered morphology.

**Figure 2.**
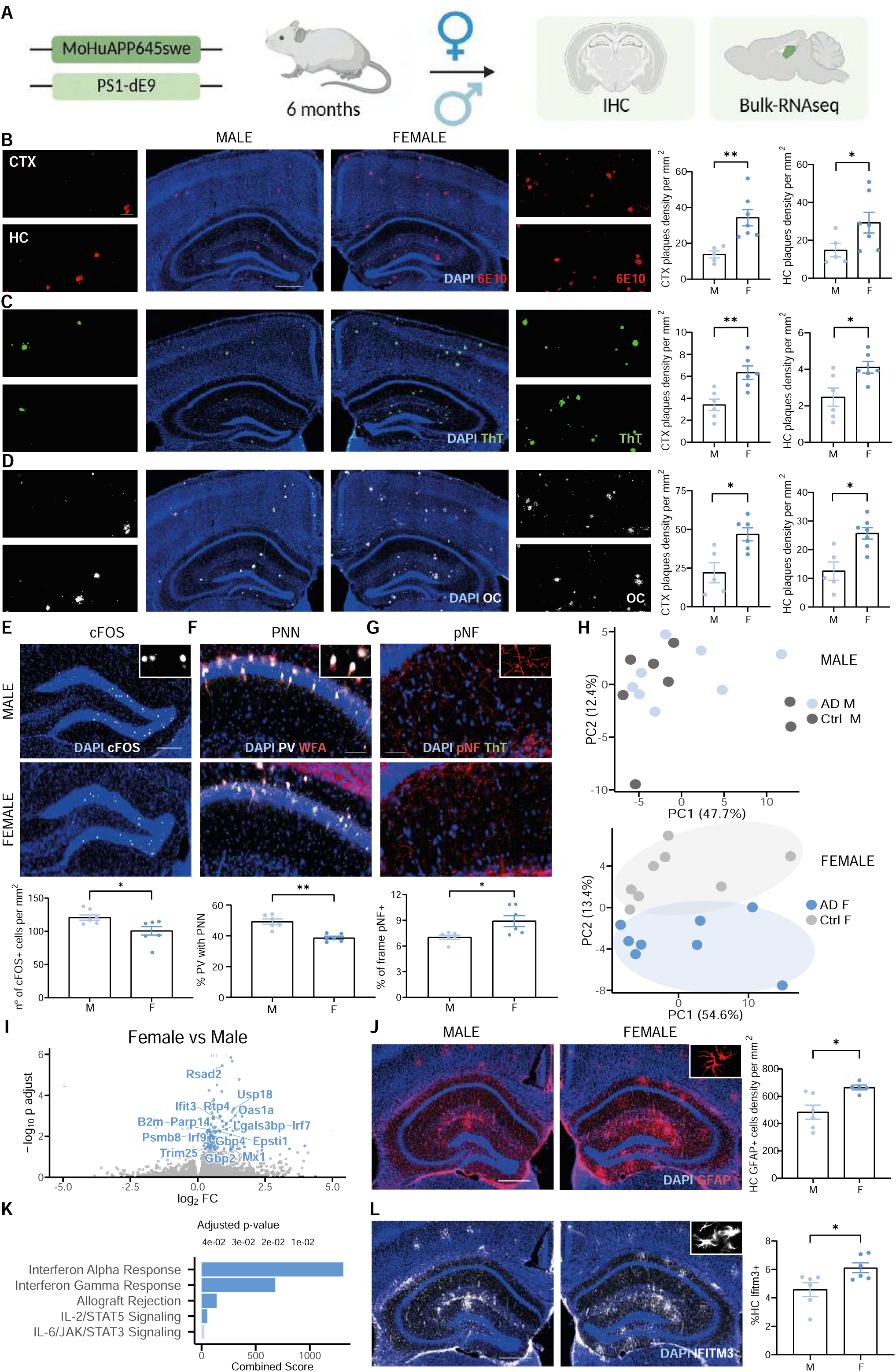
Sex differences in amyloid pathology and interferon signaling in APP/PS1 mice. (A) Schematic diagram of the APPswe/PSEN1dE9 (APP/PS1) mouse model, with experiments conducted at 6 months of age. (B) IHC with 6E10 antibody showing Aβ plaque density in cortex (CTX, \**\*P* < 0.01) and hippocampus (HC, *\*P* < 0.05) of male and female mice, with higher burden in females (quantified graphs, Mann-Whitney test). Scale bar, 500 µm; inset 100 µm. (C) Thioflavin T staining of fibrillar Aβ deposits in CTX (\**\*P* < 0.01) and HC (*\*P* < 0.05), with increased density in females (quantified graphs, Mann-Whitney test). Scale bar, 500 µm; inset 100 µm. (D) OC antibody staining of β-sheet-rich Aβ conformations in CTX and HC, showing greater signal in females (quantified graphs, *\*P* < 0.05, Mann-Whitney test). Scale bar, 500 µm; inset 100 µm. (E) c-FOS immunostaining in dentate gyrus, indicating reduced neuronal activity in female mice (quantified graphs, *\*P* < 0.05, Mann-Whitney test). Scale bar, 200 µm. (F) PNNs stained with PV/WFA in CA1, with lower proportion of PV+ interneurons in females (quantified graphs, \**\*P* < 0.01, Mann-Whitney test). Scale bar, 100 µm. (G) pNF staining for axonal damage in HC, with higher levels in females (quantified graphs, *\*P* < 0.05, Mann-Whitney test). Scale bar, 50 µm. (H) PCA of hippocampal RNA-seq data, showing greater transcriptomic divergence in female APP/PS1 mice. (I) Volcano plot of differentially expressed genes in female vs. male hippocampus, highlighting upregulated interferon alpha genes. (L) GFAP immunohistochemistry in HC, with increased cell density in females (quantified graphs, *\*P* < 0.05, Mann-Whitney test). Scale bar, 500 µm. (K) Functional enrichment analysis indicating an enrichment in the interferon alpha and gamma response pathways in female APP/PS1 mice. (L) IFITM3 immunohistochemistry in HC, with increased labeling in females (quantified graphs, *\*P* < 0.05, Mann-Whitney test). Scale bar, 500 µm. Data from 6-month-old APP/PS1 mice. IHC: n = 12 (6 females, 6 males). RNA-seq: n = 16 (8 females, 8 males). Data represent mean ± SEM.

Building on these findings, we next examined other neuropathological features associated with AD (59–63). Immunostaining for c-FOS, a marker of neuronal activity, revealed reduced expression in the dentate gyrus of female mice (Figure 2E), indicating potential functional impairment. To assess inhibitory network integrity, we quantified perineuronal nets (PNNs), specialized extracellular matrix structures that stabilize synaptic connectivity (64). The proportion of parvalbumin-positive interneurons surrounded by PNNs in the CA1 region was significantly lower in females, as visualized by *Wisteria floribunda* agglutinin (WFA) staining (Figure 2F). In parallel, we observed increased axonal damage in the female hippocampus, as evidenced by elevated phosphorylated neurofilament (pNF) staining (Figure 2G). These converging histopathological alterations suggest that female APP/PS1 mice experience more severe neuronal dysfunction and structural disruption.

To determine whether these sex-specific differences extended to the transcriptional level, we first characterized the global molecular profile of APP/PS1 mice relative to wild-type controls. Bulk RNA-Seq of hippocampal tissue revealed pronounced upregulation of transcripts related to the model’s transgenes (*App*, *Prnp*) (65) alongside microglial activation markers (*Clec7a*, *Trem2*, *Cst7, Itgax*) (66–68) (Supplemental Figure 2E and Supplemental Table 2). Functional enrichment analysis indicated significant activation of pathways including “Cholesterol Homeostasis”, “Allograft Rejection”, and “Inflammatory Response” (Supplemental Figure 2F), confirming that the APP/PS1 model exhibits the expected immune–metabolic alterations associated with AD.

Building on this baseline characterization, we next examined whether the transcriptional response to Aβ pathology differed between sexes. PCA revealed limited separation between wild-type and APP/PS1 males, whereas APP/PS1 females diverged markedly from wild-type females (Figure 2H), indicating more extensive transcriptional remodeling in females at this age (six month old). This suggests that female brains may undergo a more profound molecular response to Aβ pathology, potentially contributing to their heightened vulnerability.

These data indicate that female APP/PS1 mice exhibit greater amyloid deposition, more pronounced neurodegeneration, and broader transcriptomic alterations than males. Importantly, this model recapitulates the sex-specific vulnerability observed in human Alzheimer’s disease, making it a valuable platform for investigating the biological mechanisms underlying these differences.

### Exacerbated Neuropathology in Female APP/PS1 Mice Is Associated with Heightened Interferon Signaling

To investigate potential molecular correlates of the more pronounced AD-like histopathology observed in female APP/PS1 mice, we profiled gene expression of bulk hippocampal tissue. In line with the differences found in AD patient’s (Figure 1), differential gene expression analysis revealed a significant upregulation of inflammatory genes in females, including IFN-I-stimulated transcripts such as *Gbp4*, *Psmb8*, *Epsti1*, *Irf7*, *Irf9*, *B2m*, *Ifit3*, *Oas1a*, and *Rtp4* (Figure 2I and Supplemental Table 2).

Given the observed transcriptional signature, we examined glial activation using GFAP, a marker of astrogliosis and neuroinflammation (59, 69). Immunostaining revealed higher densities of GFAP-positive astrocytes in the hippocampi of female mice (Figure 2J), with similar increases observed in the cortex (Supplemental Figure 2G). While this suggested enhanced neuroinflammation in females, GFAP is a broad marker and does not reveal which specific inflammatory pathways are engaged. To further dissect the molecular nature of immune activation, we performed functional analysis on significantly differentially expressed genes using the “hallmark” gene sets collection (57). This revealed a selective overrepresentation of interferon-related pathways in females, including “Interferon Alpha Response” and “Interferon Gamma Response” (Figure 2K), indicating that the inflammatory profile in females is characterized by a pronounced activation of antiviral IFN-I signaling. GSEA further supported these findings, revealing a significant enrichment of the interferon alpha response gene set in APP/PS1 females (Supplemental Figure 2H).

To validate these transcriptomic findings at the protein level, we assessed the expression of IFITM3, a key effector of the interferon response. Immunohistochemistry revealed significantly increased numbers of IFITM3-positive cells in the hippocampus of females compared with males (Figure 2L), with similar results observed in the cortex (Supplemental Figure 2I).

Overall, our data indicate that the greater neurodegeneration and amyloid pathology observed in female APP/PS1 mice is accompanied by a selectively heightened interferon response, a feature that only emerged upon sex-stratified analysis, suggesting interferon-related mechanisms as a potential driver of sex-specific vulnerability in AD.

### Acute Interferon Signaling Elicits Alzheimer-Like Histopathology

Given the previously observed correlation between elevated interferon signaling and worsened neuropathological features in female APP/PS1 mice, we next examined whether interferon activation alone is sufficient to induce similar pathological changes. To test this hypothesis, we administered a single intraperitoneal dose of the viral mimetic polyinosinic:polycytidylic acid [poly(I:C), 12 mg/kg] to 3-month-old wild-type C57BL/6 mice and analyzed brain tissue 24-and 72-hours post-injection (Figure 3A).

**Figure 3.**
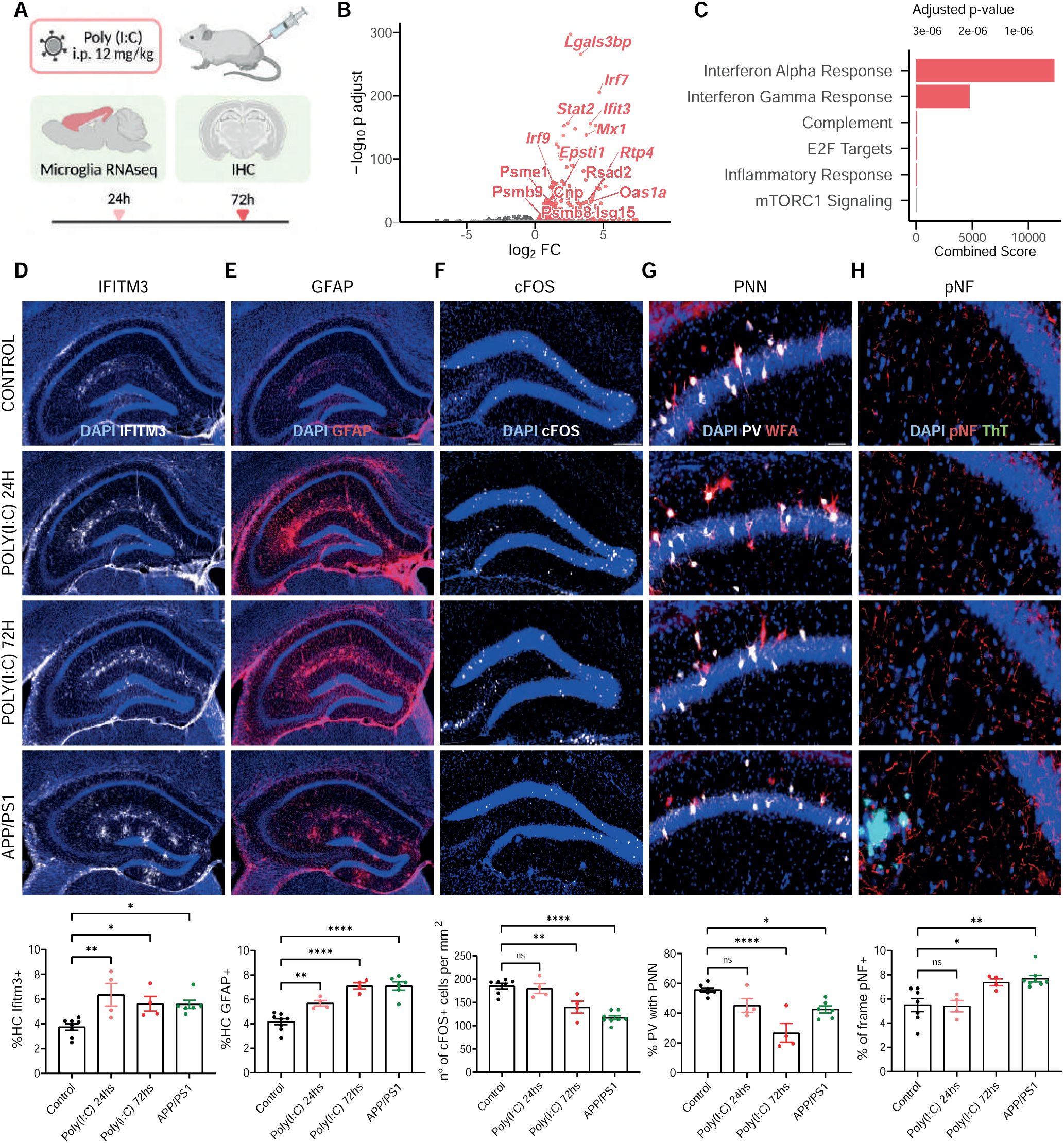
Acute interferon signaling elicits Alzheimer-like pathology. (A) Schematic of experimental design with intraperitoneal poly(I:C) administration (12 mg/kg) to 3-month-old wild-type C57BL/6 mice, analyzed at 24 and 72 hours, and compared to APP/PS1 mice using microglia RNA-sequencing and immunohistochemistry (IHC). (B) Volcano plot of differentially expressed genes in hippocampus 24 hours post-poly(I:C), highlighting upregulated interferon alpha-related genes. (C) Functional enrichment analysis showing activation of interferon alpha/gamma pathways in poly(I:C)-treated mice. (D) IFITM3 immunohistochemistry in hippocampus, with increased labeling in poly(I:C) 24h (*\*P* < 0.05), 72h (\**\*P* < 0.01), and APP/PS1 (\**\*P* < 0.01) compared to controls (quantified graphs, one-way ANOVA with Bonferroni post hoc test). Scale bar, 200 µm. (E) GFAP immunohistochemistry in hippocampus, with increased labeling in poly(I:C) 24h (\**\*P* < 0.01), 72h (\*\*\**\*P* < 0.0001), and APP/PS1 (\*\*\**\*P* < 0.0001) compared to controls (quantified graphs, one-way ANOVA with Bonferroni post hoc test). Scale bar, 200 µm. (F) c-FOS immunostaining in dentate gyrus, showing reduced neuronal activity in poly(I:C) 72h (\**\*P* < 0.01), and APP/PS1(\*\*\**\*P* < 0.0001) vs. controls (quantified graphs, one-way ANOVA with Bonferroni post hoc test). Scale bar, 200 µm. (G) PNNs stained with PV/WFA in CA1, with lower proportion of PV+ interneurons in poly(I:C) 72h (\*\*\**\*P* < 0.0001), and APP/PS1 (*\*P* < 0.05) vs. controls (quantified graphs, one-way ANOVA with Bonferroni post hoc test). Scale bar, 50 µm. (H) pNF staining for axonal damage in hippocampus, with higher levels in poly(I:C) 72h (*\*P* < 0.05), and APP/PS1 (\**\*P* < 0.01) vs. controls (quantified graphs, one-way ANOVA with Bonferroni post hoc test). Scale bar, 50 µm. Data from wild-type and APP/PS1 mice. IHC: n = 8 (control), 4 (Poly(I:C) 24h), 4 (Poly(I:C) 72h), and 6 (APP/PS1). Poly(I:C) groups included only males; other groups were sex-balanced (3 females, 3 males for APP/PS1; 4 females, 4 males for controls). RNA-seq: n = 4 (control), 3 (Poly(I:C) 24h). Data represent mean ± SEM.

First, we assessed the molecular response to poly(I:C) by profiling the transcriptome of acutely isolated cortical microglia from adult mice 24 hours after injection. PCA demonstrated clear separation between control and treated samples, indicating a robust transcriptional shift (Supplemental Figure 3A). Differential expression analysis revealed strong upregulation of interferon-related genes, including *Cnp*, *Psmb9*, *Psmb8*, *Epsti1*, *Irf7*, *Irf9*, *Ifit3*, *Oas1a*, *Rtp4*, *Stat2*, *Rsad2*, and *Isg15* (Figure 3B and Supplemental Table 3). Functional analysis confirmed significant activation of pathways involved in interferon signaling and antiviral responses, such as “Interferon Alpha Response”, “Interferon Gamma Response”, and “Complement” (Figure 3C, Supplemental Figure 3B, and Supplemental Table 3).

We next validated these transcriptional changes at the protein level by examining IFITM3 expression in relation to APP/PS1 pathology. Immunohistochemistry revealed increased hippocampal IFITM3 labeling in poly(I:C)-treated mice at 24- and 72-hours post-injection, reaching levels similar to those observed in APP/PS1 mice (Figure 3D). A similar increase was noted in the cortex (Supplemental Figure 3C), suggesting a shared interferon-related molecular signature between acute poly(I:C) stimulation and chronic APP/PS1 pathology.

Given the marked interferon response, we assessed neuroinflammatory responses by quantifying GFAP-positive astrocytes. Poly(I:C)-treated mice showed marked astrogliosis in both the hippocampus (Figure 3E) and cortex (Supplemental Figure 2D), mirroring the elevated GFAP levels observed in APP/PS1 mice. These findings indicate that poly(I:C) is sufficient to elicit glial reactivity and inflammatory signaling in the brain.

To determine whether this interferon-driven inflammatory response could also trigger structural and functional changes characteristic of AD, we next analyzed key histopathological features. At 72 hours post poly(I:C) treatment, reduced cFOS immunostaining in the dentate gyrus suggested decreased neuronal activity, mirroring the hypoactivity observed in APP/PS1 mice (Figure 3F). We further evaluated inhibitory circuit integrity by assessing PNNs. Poly(I:C) treatment reduced the proportion of parvalbumin-positive interneurons surrounded by PNNs, with a comparable decrease detected in APP/PS1 animals (Figure 3G). This was accompanied by a global reduction in WFA-labeled PNN structures (Supplemental Figure 3E), indicating disruption of extracellular matrix components critical for circuit stability. Finally, we assessed neurodegeneration-related pathology by measuring pNF accumulation. Poly(I:C) administration led to increased pNF immunoreactivity in the hippocampus, reaching levels comparable to those observed in APP/PS1 mice (Figure 3H), further supporting a link between interferon signaling and neuronal injury.

Together, these findings reveal that acute activation of the interferon pathway via viral-mimetic challenge is sufficient to elicit hallmark Alzheimer-related histopathological alterations in healthy mice. These results establish a mechanistic link between interferon responses, neuroinflammation, and neurodegeneration, supporting a causative role for interferon activation in the pathogenesis of Alzheimer’s-like features.

### Genetic Amplification of Interferon Signaling Exacerbates Alzheimer’s-Related Neuropathology

We recently showed that *Rela* targeting in microglia leads to transcriptional reprogramming towards an interferon-responsive state (Bhojwani-Cabrera et al., submitted manuscript). Taking advantage of this model and to assess the impact of chronic interferon upregulation on AD progression, we generated a gain-of-function model by crossing APP/PS1 mice with a tamoxifen-inducible, microglia-specific *Rela* knockout line (Cx3cr1::CreERT2; Rela^fl/fl^), hereafter referred to as AD hiIFN (Figure 4A). Mice were sacrificed at 6 months, allowing us to evaluate the impact of sustained microglia-driven interferon upregulation during a critical window of Alzheimer’s pathology progression.

**Figure 4.**
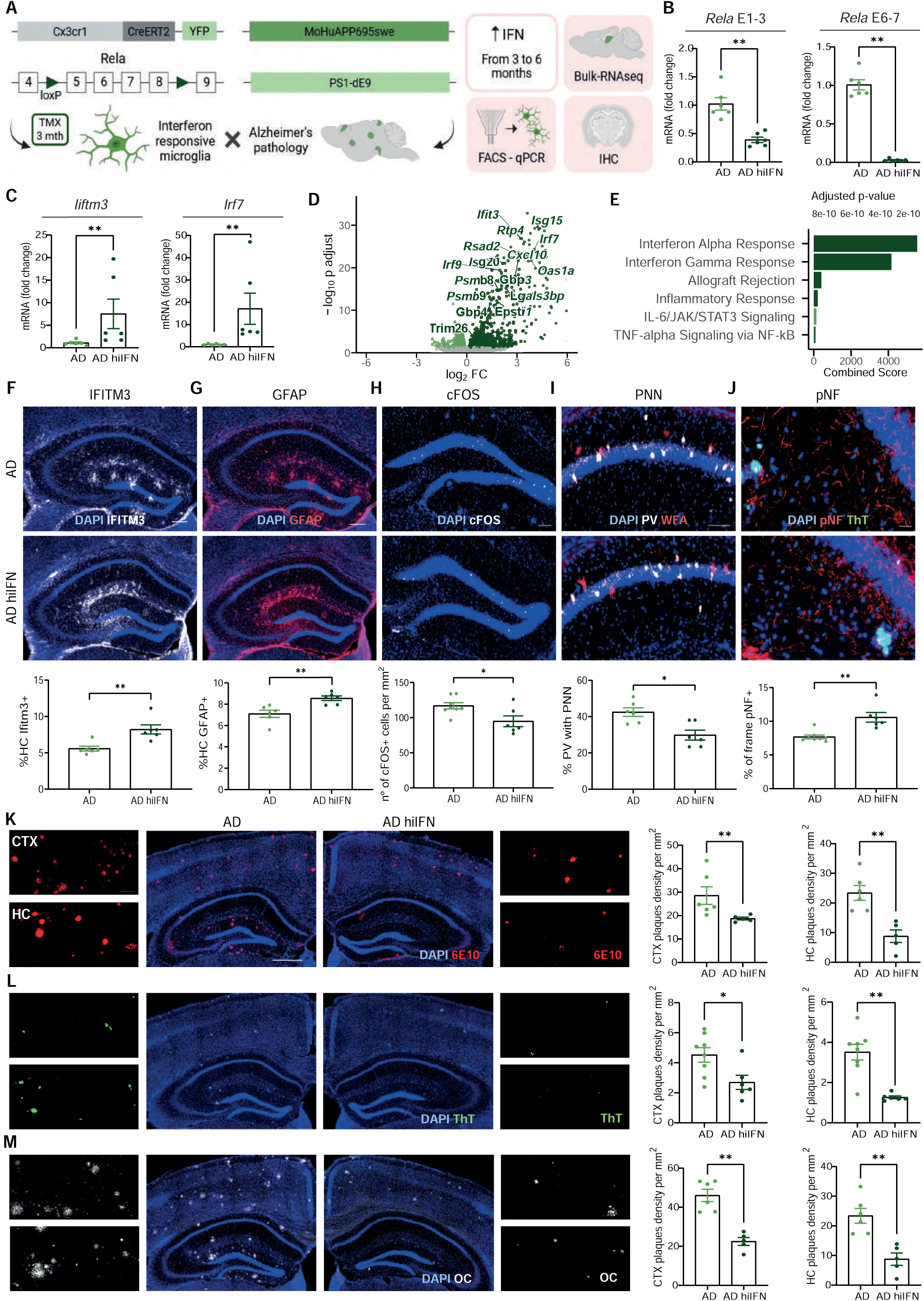
Heightened interferon signaling exacerbates Alzheimer’s pathology. (A) Schematic of APP/PS1 (AD) crossed with microglia-specific *Rela* knockout (AD hiIFN) model, with tamoxifen induction at 3 months and analysis at 6 months using qPCR, bulk-RNAseq and immunohistochemistry (IHC). (B) qPCR showing *Rela* deletion efficiency in microglia (exons 1-3 and 6-7) in AD hiIFN vs. AD (quantified graphs, \**\*P* < 0.01, Mann-Whitney test). (C) Microglia qPCR showing increased expression of interferon-related genes in AD hiIFN hippocampus (quantified graphs, \**\*P* < 0.01, Mann-Whitney test). (D) Volcano plot of differentially expressed genes in AD hiIFN vs. APP/PS1 controls, highlighting upregulated interferon alpha genes. (E) Functional enrichment analysis showing interferon alpha/gamma response pathways in AD hiIFN. (F) IFITM3 immunohistochemistry in hippocampus, with higher labeling in AD hiIFN vs. AD (quantified graphs, \**\*P* < 0.01, Mann-Whitney test). Scale bar, 200 µm. (G) GFAP immunohistochemistry for astrogliosis in hippocampus, showing increased density in AD hiIFN vs AD (quantified graphs, \**\*P* < 0.01, Mann-Whitney test). Scale bar, 200 µm. (H) c-FOS immunohistochemistry in dentate gyrus, indicating reduced neuronal activity in AD hiIFN vs. AD (quantified graphs, *\*P* < 0.05, Mann-Whitney test). Scale bar, 200 µm. (I) PNNs stained with PV/WFA in CA1, with lower proportion of PV+ interneurons in AD hiIFN vs. AD (quantified graphs, *\*P* < 0.05, Mann-Whitney test). Scale bar, 100 µm. (J) pNF staining for axonal damage in hippocampus, with higher levels in AD hiIFN vs. AD (quantified graphs, \**\*P* < 0.01, Mann-Whitney test). Scale bar, 20 µm. (K) 6E10 immunohistochemistry for Aβ plaque density, showing reduced burden in AD hiIFN vs. AD both in cortex (CTX) and hippocampus (HC) (quantified graphs, *\*P* < 0.05, Mann-Whitney test). Scale bar, 500 µm; inset 100 µm. (L) Thioflavin T staining for fibrillar Aβ deposits, with lower density in AD hiIFN vs. AD in CTX (*\*P* < 0.05) and HC (\**\*P* < 0.01), (quantified graphs, Mann-Whitney test). Scale bar, 500 µm; inset 100 µm. (M) OC staining for β-sheet-rich Aβ conformations, with reduced signal in AD hiIFN vs. AD in CTX and HC, (quantified graphs, \**\*P* < 0.01, Mann-Whitney test). Scale bar, 500 µm; inset 100 µm. Data from 6-month-old AD hiIFN and APP/PS1 mice. IHC: n = 6 per group (3 females, 3 males). RNA-seq: n = 8 per group (4 females, 4 males). Data represent mean ± SEM.

Efficient deletion of *Rela* in microglia was confirmed by qPCR performed on fluorescence-activated cell sorted (FACS) microglia, showing significant reduction in exons 1–3 and 6–7 expression, with a more pronounced decrease in exons 6–7 (Figure 4B). This genetic manipulation resulted in increased expression of interferon-related genes, including *Irf7*, and *Ifitm3* (Figure 4C).

Bulk RNA sequencing of hippocampal tissue revealed robust upregulation of interferon-associated genes such as *Gbp4*, *Psmb9*, *Psmb8*, *Epsti1*, *Irf7*, *Irf9*, *Ifit3*, *Oas1a*, *Rtp4*, *Rsad2*, and *Isg15* in AD hiIFN mice compared to AD controls (Figure 4D and Supplemental Table 4). PCA confirmed clear transcriptional separation between genotypes, supporting a distinct molecular signature induced by chronic interferon activation (Supplemental Figure 4A). Gene ontology enrichment analysis further highlighted significant overrepresentation of interferon and viral defense pathways, including “Interferon Alpha Response” and “Interferon Gamma Response” (Figure 4E and Supplemental Table 4). Gene set enrichment analysis using the “hallmark” gene set collection (57) confirmed enrichment of “Interferon Alpha Response” pathway in AD hiIFN mice (Supplemental Figure 4B).

To determine whether this transcriptional program was influenced by sex, we compared interferon responses across male and female mice in the AD hiIFN model. Analysis of the top 15 significantly upregulated interferon-stimulated genes AD hiIFN revealed that both sexes reached comparable levels of interferon-related transcripts (Supplemental Figure 4C). Based on this observation, both male and female animals were included in the analysis. Protein-level validation by IFITM3 immunohistochemistry confirmed elevated interferon activity in AD hiIFN mice, with increased labeling of IFITM3 in the hippocampus (Figure 4F) and cortex (Supplemental Figure 4D).

Having confirmed sustained interferon activation at both the transcriptomic and protein levels we next assessed the consequences on key histopathological features. GFAP immunostaining revealed markedly exacerbated astrogliosis in AD hiIFN mice, with higher GFAP labeling in hippocampus (Figure 4G) and, nearly to significance, in cortex (Supplemental Figure 4E). In the dentate gyrus, AD hiIFN mice showed a more pronounced reduction of c-FOS expression than AD mice, suggesting further impairment of neuronal activity (Figure 4H). Structural alterations in inhibitory interneuron networks were also apparent, with a decreased proportion of parvalbumin-positive interneurons enwrapped by perineuronal nets in CA1 (Figure 4I) and a global reduction in WFA-labeled structures that approached significance (Supplemental Figure 4F). Finally, axonal integrity was impaired, shown by increased phosphorylated neurofilament staining in hippocampus (Figure 4J).

Interestingly, despite these exacerbated signs of neurodegeneration and gliosis, amyloid plaque burden was reduced in AD hiIFN mice. Quantification with 6E10 (Figure 4K), Thioflavin T (Figure 4L), and OC antibodies (Figure 4M) demonstrated lower plaque density in both cortex and hippocampus, with largely comparable plaque size distribution (Supplemental Figure 4, G-I).

These data identify IFN-I signaling as a disease-modifying mechanism in AD. Chronic upregulation of the IFN-I axis exacerbates neuroinflammation and neurodegeneration, uncoupling plaque load from neuronal damage and underscoring a pathogenic role for interferon-driven processes in disease progression. These findings highlight IFN-I signaling as a potential therapeutic target in AD.

### Modulation of Interferon Signaling Via STING Antagonism Ameliorates Alzheimer’s-Associated Histopathology

Having demonstrated that elevated IFN-I signaling worsens AD pathology, we next explored whether attenuating this pathway could yield neuroprotective effects. To address this, APP/PS1 mice received intraperitoneal injections of C-176, a selective STING antagonist that prevents Cys91 palmitoylation and thereby blocks downstream interferon production (70), in alternate days from 3 to 6 months of age, matching the treatment window used in the IFN-I gain-of-function model (Figure 5A). These animals are hereafter termed AD loIFN.

**Figure 5.**
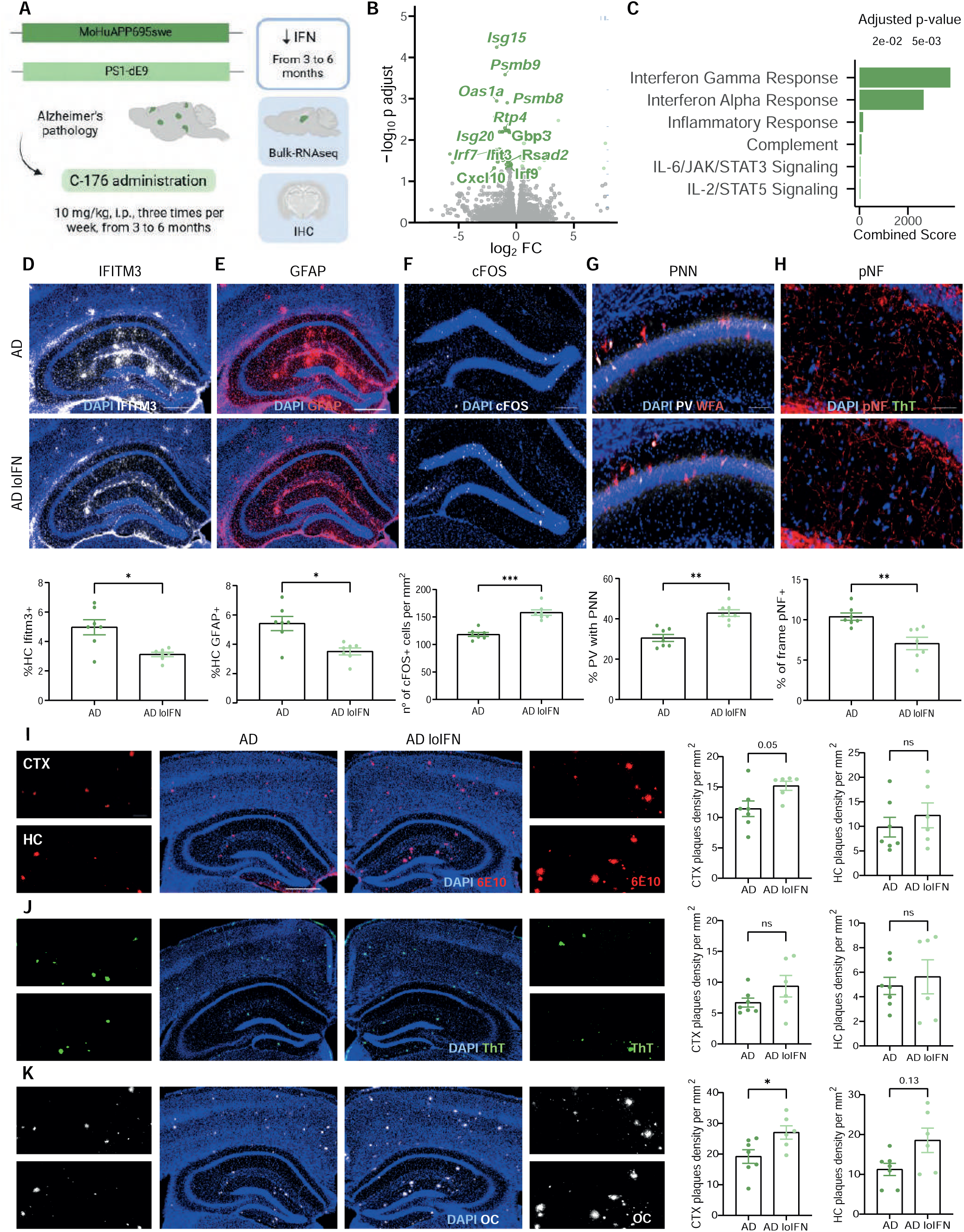
STING inhibition alleviates Alzheimer’s pathology in APP/PS1 mice. (A) Schematic of experimental design with STING inhibitor C-176 (10 mg/kg) administration to 3-month-old APP/PS1 mice (AD loIFN), treated until 6 months, analyzed using bulk RNA-seq and immunohistochemistry (IHC). (B) Volcano plot of differentially expressed genes in hippocampus of treated vs untreated females, highlighting downregulated interferon alpha genes. (C) Functional enrichment analysis showing reduced interferon alpha/gamma response pathways in untreated vs. treated females. (D) IFITM3 immunohistochemistry in hippocampus, with lower labeling in treated AD loIFN vs. untreated AD mice (quantified graphs, *\*P* < 0.05, Mann-Whitney test). Scale bar, 500 µm. (E) GFAP immunohistochemistry for astrogliosis in hippocampus, showing decreased density in treated vs. untreated mice (quantified graphs, *\*P* < 0.05, Mann-Whitney test). Scale bar, 500 µm. (F) c-FOS immunohistochemistry in dentate gyrus, indicating increased neuronal activity in treated vs. untreated mice (quantified graphs, \*\**\*P* < 0.001, Mann-Whitney test). Scale bar, 200 µm. (G) PNNs stained with PV/WFA in CA1, with higher proportion of PV+ interneurons in treated AD loIFN vs. untreated AD mice (quantified graphs, \**\*P* < 0.01, Mann-Whitney test). Scale bar, 100 µm. (H) pNF staining for axonal damage in hippocampus, with lower levels in treated vs. untreated mice (quantified graphs, \**\*P* < 0.01, Mann-Whitney test). Scale bar, 50 µm. (I) 6E10 immunohistochemistry for Aβ plaque density, showing non-significant changes in AD loiIFN vs. AD both in cortex (CTX) and hippocampus (HC) (quantified graphs, ns, *P* > 0.05, Mann-Whitney test). Scale bar, 500 µm; inset 100 µm. (J) Thioflavin T staining for fibrillar Aβ deposits, with no changes in treated vs. untreated mice both in CTX and HC quantified graphs, ns, *P* > 0.05, Mann-Whitney test). Scale bar, 500 µm; inset 100 µm. (K) OC staining for β-sheet-rich Aβ conformations, with increased signal in AD loIFN vs. AD in CTX (*\*P* < 0.05) and non-significant changes in HC (ns, *P* > 0.05) (quantified graphs, Mann-Whitney test). Scale bar, 500 µm; inset 100 µm. Data from 6-month-old AD loIFN and APP/PS1 mice. IHC: n = 6 per group (3 females, 3 males). RNA-seq: n = 8 AD (females), 4 AD loIFN (females). Data represent mean ± SEM.

To validate the immunomodulatory effect of C-176, we first treated BV2 microglial cultures with poly(I:C) in the presence or absence of the compound. C-176 reduced expression of *Irf7*, *Ifitm3*, and *Stat2* while increasing levels of the anti-inflammatory cytokine *Il-10*, supporting its efficacy in suppressing interferon signaling in vitro (Supplemental Figure 5A).

We next evaluated the transcriptional effects of targeting IFN-I signaling *in vivo*. Given the sex differences observed in IFN-I activation across previous experiments, we first assessed whether the transcriptional effects of C-176 were comparable between male and female APP/PS1 mice. To this end, we assessed the top 15 significantly downregulated interferon-stimulated genes in AD loIFN model. This analysis revealed a robust suppression of interferon-related transcripts exclusively in C-176–treated females, while gene expression in males remained largely unchanged (Supplemental Figure 5B). These findings suggest that the efficacy of interferon pathway inhibition is sex-dependent, likely due to the already low interferon tone observed in untreated AD males. Based on these results, transcriptomic analyses were focused on female mice, in whom C-176 produced a measurable molecular response.

Accordingly, differential gene expression analysis of hippocampal tissue from treated versus untreated female APP/PS1 mice revealed significant downregulation of key interferon-stimulated genes, including *Psmb9*, *Psmb8*, *Irf7*, *Irf9*, *Ifit3*, *Oas1a*, *Rtp4 Rsad2*, and *Isg15* (Figure 5B and Supplemental Table 5). Functional analysis indicated a loss of interferon-and viral defense–associated “hallmark” gene sets in the treated group, including “Interferon Gamma Response” and “Interferon Alpha Response” (Figure 5C and Supplemental Table 5). GSEA further confirmed significant negative enrichment of the “Interferon Alpha Response” pathway elicited by C-176 treatment (Supplemental Figure 5C). Analysis of male-specific datasets confirmed a minimal transcriptional response to C-176, with no significant changes in interferon-related genes or pathways detected (Supplemental Figure 5, D and E).

Consistent with overall transcriptional changes, protein-level validation by immunohistochemistry showed reduced numbers of IFITM3-positive cells in the hippocampus (Figure 5D) and cortex (Supplemental Figure 5F) of treated mice.

Critically, these transcriptional and protein-level effects coincided with improvements in AD-related histopathological features. GFAP immunostaining revealed a significant reduction in astrogliosis in hippocampus (Figure 5E) with a similar trend observed in cortex (Supplemental Figure 5G). Neuronal activity, measured by c-FOS expression in the dentate gyrus, was partially restored following C-176 treatment (Figure 5F). The integrity of inhibitory circuits was also preserved, as reflected by an increased proportion of parvalbumin-positive interneurons surrounded by PNNs (Figure 5G) and a concomitant rise in WFA-labeled PNN structures (Supplemental Figure 5H). Axonal integrity was likewise improved, as evidenced by a decrease in pNF staining in the hippocampus (Figure 5H).

Notably, despite these improvements in neuroinflammation and neuronal preservation, amyloid pathology remained largely unchanged. Immunostaining with 6E10 revealed no significant differences in plaque burden in the hippocampus, with only a modest, non-significant increase in the cortex (Figure 5I). Thioflavin T staining confirmed the absence of changes in plaque load across brain regions, whereas OC immunostaining showed no significant difference in the hippocampus but a significant increase in plaque burden in the cortex of treated mice (Figure 5, J and K). Importantly, across all three markers, plaque size showed a consistent trend toward reduction in treated animals (Supplemental Figure 5, I–K), suggesting that IFN-I pathway inhibition may preferentially impact plaque maturation and compaction rather than initial deposition.

In summary, pharmacological inhibition of IFN-I signaling via STING antagonism mitigates key AD-related histopathological features, dampening neuroinflammation and preserving neuronal integrity independently of plaque load and pointing to IFN-I pathway suppression as a promising therapeutic approach for modifying disease trajectory.

### Targeting Type I Interferon Signaling Preserves Cognitive Function in Female APP/PS1 Mice

Given our findings that heightened IFN-I signaling worsens Alzheimer pathology, while its pharmacological inhibition alleviates key histological hallmarks, we next investigated whether modulation of IFN-I activity also influences cognitive performance (Figure 6A). Specifically, we aimed to establish whether IFN-I signaling serves as a functional mediator linking neuroimmune dysregulation to behavioral impairments in AD.

**Figure 6.**
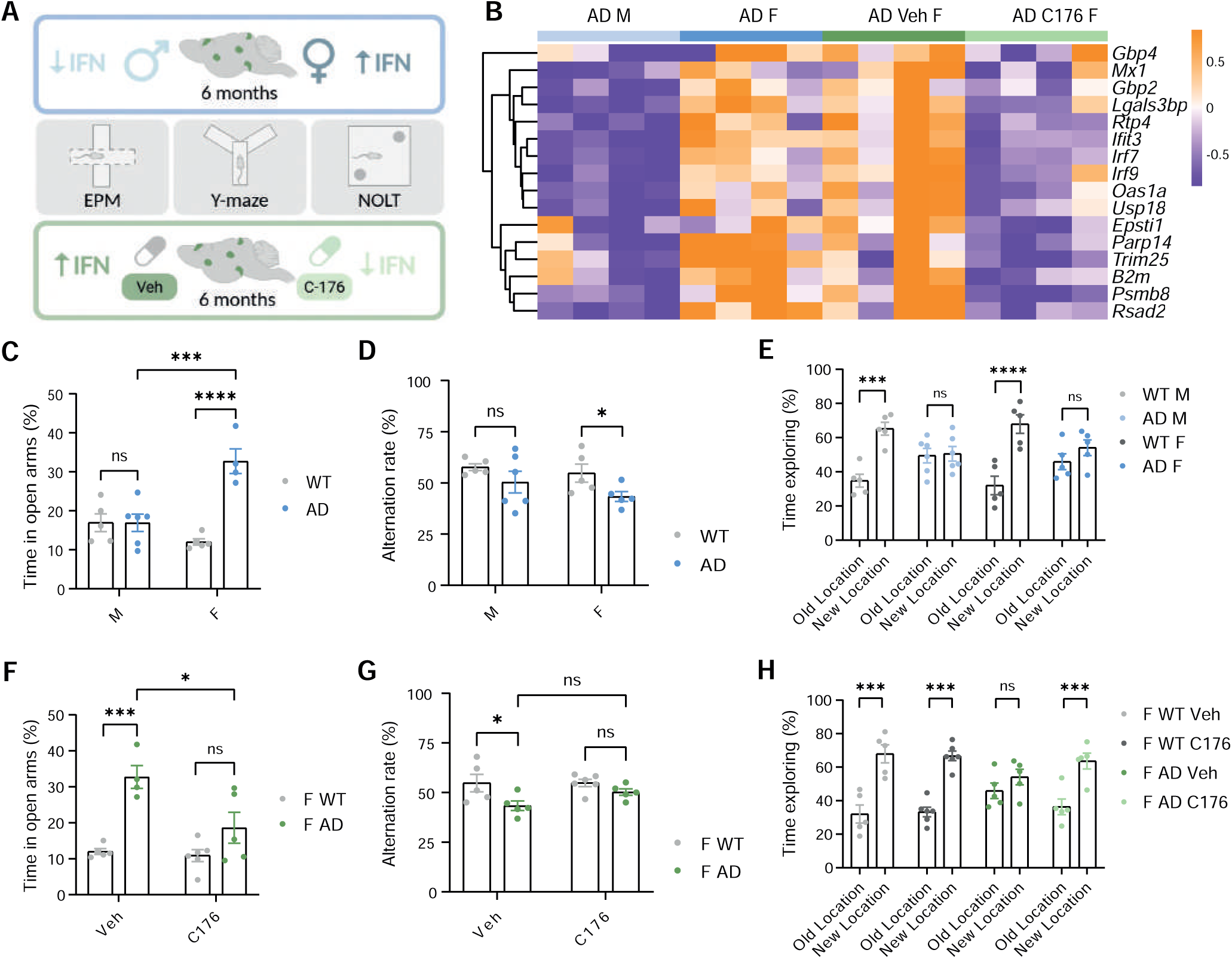
Targeting Interferon Signaling Preserves Cognitive Performance in Female APP/PS1 Mice. (A) Schematic of experimental design for cognitive testing in male and female APP/PS1 mice (AD) and female APP/PS1 mice treated with C-176 (AD loIFN) from 3 to 6 months. (B) Heatmap of the top 15 upregulated interferon-alpha– related genes in hippocampal tissue from female AD mice, showing elevated expression compared to both male AD mice and C-176–treated female AD mice. (C) Elevated Plus Maze (EPM) results, showing increased time in open arms in female AD mice compared to WT females (\*\*\**\*P* < 0.0001), with no differences in males (ns, *P* > 0.05) (quantified graphs, \*\**\*P* < 0.001, one-way ANOVA with Bonferroni post hoc test). (D) Y-Maze spontaneous alternation, revealing reduced alternation in female AD mice compared to WT (*\*P* < 0.05), with no differences in males (ns, *P* > 0.05) (quantified graphs, one-way ANOVA with Bonferroni post hoc test). (E) Novel Object Location Test (NOLT) results, indicating impaired spatial memory in both male and female AD mice compared to WT (quantified graphs, \*\**\*P* < 0.001, \*\*\**\*P* < 0.0001, ns *P* > 0.05, one-way ANOVA with Bonferroni post hoc test). (F) EPM in AD C-176-treated females, indicating normalized time in open arms compared to treated WT females (ns, *P* > 0.05) and demonstrating cognitive improve compared to untreated AD females (*\*P* < 0.05) (quantified graphs, \*\**\*P* < 0.001, one-way ANOVA with Bonferroni post hoc test). (G) Y-maze in AD C-176-treated females, indicating normalized time in open arms compared to treated WT females (ns, *P* > 0.05) (quantified graphs, *\*P* < 0.05, one-way ANOVA with Bonferroni post hoc test). (H) NOLT in C-176-treated female AD mice (AD loIFN), showing restored discrimination of novel object location compared to untreated AD females (quantified graphs, \*\**\*P* < 0.001, ns *P* > 0.05, one-way ANOVA with Bonferroni post hoc test). Data from 6-month-old WT, AD, and AD loIFN mice. n = 5 (male WT), 6 (male AD), 5 (female WT), 5 (female AD), 5 (female WT + vehicle), 5 (female WT + C-176), 5 (female AD + vehicle), and 5 (female AD + C-176). Data represent mean ± SEM.

We first confirmed group-specific IFN-I activity by transcriptomic profiling of hippocampal tissue. Comparison between female and male APP/PS1 mice revealed selective upregulation of ISGs in females, including *Gbp4*, *Psbm9*, *Psbm8*, *Epsti1*, *Irf7*, *Irf9*, *Ifit3, Oas1a*, *Rtp4*, and *Rsad2*. These transcriptional changes were all attenuated by C-176 treatment in females (Figure 6B), indicating sex-dependent regulation of interferon signaling and its pharmacological modulation.

Behavioral testing revealed domain-specific impairments in APP/PS1 mice, with a clear sex bias emerging across tasks of increasing complexity. In the elevated plus maze (EPM), no significant differences were observed between APP/PS1 males and their wild-type counterparts. In contrast, APP/PS1 females spent significantly more time in the open arms compared to wild-type females (Figure 6C), consistent with altered anxiety-like behavior. Similarly, Y-maze testing revealed impaired spontaneous alternation in females - indicative of working memory deficits - while male APP/PS1 mice again performed comparably to controls (Figure 6D). However, in the novel object location test (NOLT), which places higher demands on spatial memory, both male and female transgenic mice failed to recognize the novel location (Figure 6E). These findings parallel the sex-specific pattern of IFN-I activation observed at the molecular level and support a functional link between interferon dysregulation and cognitive decline.

To test this hypothesis directly, we evaluated whether suppression of IFN-I signaling could ameliorate cognitive impairments in female APP/PS1 mice. Consistent with transcriptomic evidence of ISG downregulation in the hippocampus (Figure 6B), C-176 treatment led to behavioral improvements across multiple domains. In the EPM, treated mice spent significantly less time in the open arms compared to untreated APP/PS1 females, resulting in anxiety-like behavior levels comparable to those observed in control mice (Figure 6F). In the Y-maze, C-176–treated females showed improved spontaneous alternation, reaching performance levels comparable to wild-type controls and indicative of restored working memory (Figure 6G). Finally, in the more demanding NOLT, treated mice successfully discriminated the novel location, in contrast to their untreated counterparts (Figure 6H), suggesting recovery of spatial memory function.

Notably, performance in the Open Field test was unaffected by genotype or treatment, indicating no major alterations in locomotor activity or basal anxiety (Supplemental Figure 6, A and E). Similarly, no significant differences were detected in the T-maze (Supplemental Figure 6, B and F), contextual fear memory (Supplemental Figure 6, C and G) or cued fear memory performance (Supplemental Figure 6, D and H), suggesting that not all cognitive domains are equally influenced by interferon signaling.

Together, these data demonstrate that interferon signaling not only shapes molecular and histopathological features of AD but also contributes to cognitive dysfunction. Importantly, the ability of STING antagonism to reverse behavioral deficits highlights IFN-I modulation as a promising avenue for therapeutic intervention aimed at preserving cognitive function in AD.

## Discussion

Alzheimer’s disease (AD) affects women disproportionately, both in prevalence and progression rate (1, 3, 71–73). While demographic factors such as increased lifespan contribute to this disparity, emerging evidence suggests that biological sex fundamentally shapes the neurobiological response to AD pathology (12, 74). In this study, we show that interferon signaling is selectively and robustly upregulated in the brains of female AD patients and that this sex bias in immune activation corresponds with heightened vulnerability to neurodegeneration in animal models. Through targeted manipulation of the interferon axis *in vivo*, we establish a causal relationship between interferon activity and hallmark features of AD pathology, including neuronal dysfunction, gliosis, and cognitive decline.

Our transcriptomic analyses of the parahippocampal gyrus from AD patients revealed significant enrichment of type I interferon response in females, along with increased type II interferon. These pathways, well-characterized mediators of antiviral immunity and inflammation, have been increasingly implicated in neurodegenerative contexts (75–85). This observation builds on prior reports of sex differences in peripheral immune responses (86–91) and extends them to the human brain, suggesting that female-specific amplification of innate immunity may underlie increased AD susceptibility (92–97).

To explore the functional relevance of these observations, we used the APP/PS1 mouse model of amyloidosis. Female transgenic mice exhibited more severe neuropathology than males, including greater amyloid burden, increased glial activation, and enhanced neuronal and axonal pathology. These histopathological findings were mirrored by transcriptomic alterations, with female hippocampi displaying stronger upregulation of interferon-stimulated genes and antiviral defense pathways. While sex differences in AD models have been reported before (98–102), our work provides a mechanistic framework linking female-biased IFN-I activation to downstream neuropathological consequences.

Importantly, inducing IFN-I signaling in otherwise healthy wild-type mice - via systemic administration of the viral mimetic poly(I:C) - was sufficient to reproduce several AD-like features (103–105), including reduced neuronal activity (cFOS) (104, 105), degradation of perineuronal nets (PNNs) (106), astrogliosis (107), and axonal damage (108). These findings align with prior demonstrations that chronic IFN-I impairs neuronal function and plasticity; for instance, Baruch et al. showed that IFN-I drives synaptic loss and cognitive decline in aging mice, while Blank et al. reported suppression of hippocampal neurogenesis and long-term potentiation by IFN-I (109–113). Our observation that both cFOS-positive neurons and PNNs decline following IFN-I activation supports the idea that innate immune overactivation directly disrupts neuronal circuits (81, 109, 114–116). These results extend the role of IFN-I in the CNS beyond antiviral defense, positioning it as a key regulator of neural integrity.

The use of poly(I:C) to mimic viral activation also intersects with the long-standing “infectious hypothesis” of AD, which posits that the disease may be driven, at least in part, by host immune responses to chronic or latent infections (30, 34, 111). Several studies have identified herpesvirus DNA in postmortem AD brains (117–120), and recent transcriptomic analyses have revealed molecular signatures of past viral exposure, including SARS CoV 2–related signals (121–124). Although causality remains unresolved, our data show that even sterile mimetics of viral RNA can trigger key aspects of AD pathology, reinforcing the notion that persistent immune stimulation can act as a primary driver of neurodegeneration.

In this context, it is noteworthy that population-based studies have found associations between vaccination against pathogens such as influenza and herpes zoster, and a reduced incidence of dementia, including AD (125–129). Although mechanisms remain speculative, one hypothesis is that vaccines reduce the frequency or severity of viral reactivation and associated IFN-I bursts (115, 125, 130). Alternatively, they may establish a more regulated immune tone less prone to maladaptive activation (75, 76, 128). Our data provides mechanistic support for these epidemiological associations, suggesting that modulation of IFN-I activity may mediate the protective effects of immunization.

To further assess the consequences of sustained IFN-I activation, we developed a genetic gain-of-function model in which deletion of *Rela* in microglia reprogrammed these cells toward an IFN-I-responsive phenotype (Bhojwani-Cabrera et al., submitted manuscript). In that study, *Rela* knockout mice with heightened IFN-I pathway activation exhibited cognitive impairment, consistent with previous reports linking increased interferon signaling to cognitive decline (80, 84, 110, 113, 131, 132). When crossed with APP/PS1 mice, this manipulation led to exacerbated neuroinflammation, axonal pathology, and reduced neuronal activity. Interestingly, amyloid plaque burden was reduced in these animals, indicating that IFN-I signaling can drive neurotoxicity through mechanisms that are not strictly dependent on plaque accumulation (80, 81, 85). This reflects a broader conceptual shift in the field - from viewing AD primarily as a disorder of protein accumulation to recognizing the role of maladaptive immune responses (133–135). Our findings parallel results from microglia-specific *C9orf72* knockout mice, which also showed robust IFN-I signature accompanied by synaptic loss and reduced amyloid deposition (136). However, the case of IFITM3 - a key IFN-I effector - underscores the complexity: although it enhances γ-secretase activity and amyloidogenesis (137), its upregulation in *Rela* cKO brains coincided with fewer plaques, highlighting that its effect may depend on the overall inflammatory context. Together, these observations suggest that the neurotoxic impact of IFN-I responses may outweigh their influence on plaque dynamics.

A key upstream trigger of IFN-I signaling in AD is activation of the cGAS– STING pathway, which senses cytosolic DNA such as mitochondrial DNA released by cellular stress - a process exacerbated near amyloid plaques (138–141). This pathway is aberrantly activated in both AD models and human brain tissue, promoting IFN-I responses and neuroinflammation (114, 142–145). Notably, our transcriptomic analyses revealed increased p53 pathway activity in AD females compared to males, a relevant finding given that p53 can amplify cGAS–STING signaling in response to DNA damage– associated stress (146, 147). Consistent with these findings, we targeted STING using C-176 in APP/PS1 mice. This treatment reduced IFN-I-stimulated gene expression, dampened gliosis, preserved neuronal activity and axonal integrity, and improved cognitive performance - particularly in females with higher baseline IFN-I levels. Notably, these improvements occurred despite minimal changes in plaque burden, reinforcing the idea that cGAS–STING–driven IFN-I responses contribute to neurodegeneration independently of amyloid accumulation.

This dissociation between plaque deposition and clinical outcome mirrors observations in “resilient” individuals - older adults with high amyloid load who remain cognitively intact (148, 149). Postmortem analyses of such individuals often reveal low glial activation and minimal inflammation (150, 151). Similarly, in C-176 treated mice,IFN-I signaling was suppressed and cognitive function preserved despite persistent plaques. These data strengthen the argument that the immune system - especially the IFN-I axis - is a key determinant of whether amyloidosis remains subclinical or progresses to overt neurodegeneration.

Although these results nominate the IFN-I axis as a compelling therapeutic target, translation to the clinic demands scrutiny. C-176 has demonstrated efficacy in modulating IFN-I signaling *in vivo* in murine models, yet its activity is confined to the murine STING isoform, with no evidence supporting efficacy against the human variant (70, 152). By contrast, compounds such as H-151 exhibit greater selectivity toward human STING and represent more promising candidates for clinical development (70). Importantly, preclinical data in murine models support the blood–brain barrier permeability and CNS penetrance of these inhibitors, providing a promising pharmacokinetic basis for further investigation (114, 153–155). Nonetheless, their pharmacokinetic properties in humans remain to be characterized. Although IFN-I plays an essential role in host defense and tumor surveillance, and systemic inhibition carries potential risks (116, 130, 156, 157), several IFN-I-targeting agents - such as JAK inhibitors and anti IFNAR antibodies - are already in clinical use for autoimmune disorders, providing a strong foundation for repurposing (158–160). Patient stratification based on peripheral or cerebrospinal biomarkers of IFN-I activity could further refine candidate selection and enhance therapeutic precision.

These considerations underscore the need for precision medicine approaches that account for interindividual variability in immune tone and response. Notably, our findings emphasize the importance of sex as a biological variable in both disease progression and therapeutic efficacy. Historically, females have been underrepresented in neuroscience research, and many animal models fail to account for sex-based differences (11, 21, 22, 24). Here, the inclusion of both sexes revealed critical differences in pathology and a female-specific therapeutic response to C-176 that would have been missed in a unisex design. Moreover, the observation that epithelial–mesenchymal transition emerged as the dominant pathway in males, whereas interferon alpha response predominated in females, highlights fundamentally different disease-associated programs that may each warrant tailored intervention. Future studies should address the potential pathogenic role of epithelial–mesenchymal transition in male AD, as understanding these sex-specific signatures will be essential for developing stratified therapeutic strategies in which sex and immune phenotype guide clinical decision-making.

In conclusion, our study reveals that IFN-I signaling is selectively elevated in the female AD brain and that this pathway actively contributes to neuronal and cognitive dysfunction, independently of amyloid burden. By integrating human transcriptomics with mechanistic animal models, we identify IFN-I as a central, sex-biased effector of AD pathogenesis. These findings suggest that modulating IFN-I responses may offer therapeutic benefit, particularly in patient populations characterized by heightened innate immune activation. Future research should aim to identify biomarkers of IFN-I tone, evaluate the safety of long-term IFN-I modulation, and determine whether early intervention in IFN-I-dominant disease trajectories can prevent the transition from asymptomatic amyloidosis to neurodegenerative decline.

## Methods

### Sex as a biological variable

Both male and female mice were included in all experimental groups unless otherwise stated, and sex was a central consideration in the study design. Experimental cohorts were balanced by sex, and all transcriptomic, histological, and behavioral analyses were stratified accordingly. This allowed for direct comparison of AD–related phenotypes and immune signaling between male and female animals. Similarly, human transcriptomic datasets from the The Mount Sinai Brain Bank (MSBB) study (56) included 66 AD patients (34 females and 32 males) and 42 age-matched healthy controls (19 females and 23 males), and differential expression analyses were performed to identify sex-specific molecular signatures. Collectively, these approaches ensured that sex was rigorously integrated as a biological variable throughout the study.

### Mice

APP/PS1 (65) and Cx3cr1^CreERT2^ (161) mice are available at public repositories (RRID:MMRRC_034832-JAX and RRID:IMSR_JAX:021160, respectively). RelA^fl/fl^ mice which harbor loxP sites flanking exons 5-8 of the *Rela* gene, have been previously described (MGI:3775205) (162) and were kindly provided by Dr. Angel Barco (Instituto de Neurociencias). Cx3cr1^CreERT2^ mice were crossed with RelA^fl/fl^ mice (162) resulting in the line referred as Cx3cr1^CreERT2^-Rela^fl/fl^. These mice were crossed with APP/PS1 mice to generate Cx3cr1^CreERT2^-Rela^fl/fl^-APP/PS1. In the experiments involving tamoxifen-induced knockouts, we used tamoxifen-treated Rela^fl/fl^-APP/PS1-littermates as controls. All transgenic lines were maintained on a C57BL/6J genetic background.

### BV2 microglial cell line

Mouse microglial BV2 cells were cultured in high-glucose Dulbecco’s modified Eagle’s medium (DMEM) (Thermo Fisher Scientific, 41965047) supplemented with 10% heat-inactivated fetal bovine serum (FBS) (Thermo Fisher Scientific, 10270106) and 1% penicillin-streptomycin (Sigma-Aldrich, P4333). Cells were maintained at 37°C in a humidified atmosphere with 5% CO₂ and plated at a density of 3 × 10⁵ cells per well in 6-well plates for stimulation assays.

### *In vivo* animal treatments

According to each experiment, animals were treated following ethical guides as described previously. To induce Cre recombination in Cx3cr1^CreERT2^ mice, tamoxifen (TMX) (Sigma-Aldrich, T5648) was dissolved at 25 mg/mL in corn oil (Sigma-Aldrich, C8267) by heating to 50°C for 1 hour under light-protected conditions. Three-month-old mice received two intraperitoneal injections of tamoxifen (75 mg/kg) on alternate days. All downstream experiments involving TMX were conducted at least one month after TMX administration. Polyinosinic:polycytidylic acid high molecular weight (poly(I:C)) (Sigma-Aldrich, P1530-25MG) was prepared in sterile, endotoxin-free water and administered intraperitoneally at a dose of 12 mg/kg. Mice were sacrificed at 24- or 72-hours post-injection for transcriptomic and histological analyses. The STING inhibitor C-176 (MedChemExpress, HY-112906) was prepared at 1.5 mg/mL in a corn oil:ethanol (90:10) solution. Mice received intraperitoneal injections of 10 mg/kg on alternate days from 3 to 6 months of age, corresponding to the window of APP/PS1 pathology development.

### Immunohistochemistry and image analysis

Mice were deeply anesthetized and transcardially perfused with PBS, followed by 4% paraformaldehyde (PFA) (PFA, w/v, dissolved in 0.1 M phosphate buffer, pH 7.4). Brains were post-fixed overnight and cryoprotected in 30% (w/v) sucrose for 48h. Coronal sections (50 µm) were obtained using a cryotome and stored in antifreeze solution (30% ethylene glycol, 30% glycerol, 30% distilled water, 10% PBS) at −20°C until processing. Briefly, free-floating sections were washed 3 times in PBS (5 min each) and blocked in 5% Newborn calf serum (NCS) (Sigma-Aldrich, N4762) with 0.3% Triton X-100 (Sigma-Aldrich, T8787), followed by overnight incubation at 4°C with primary antibodies. The following primary antibodies were used: IFITM3 (Proteintech, 11714-1-AP, 1:500), 6E10 (anti-β-Amyloid, 1–16) (BioLegend, 803001, 1:1000), c-FOS (Synaptic Systems, 226308, 1:1000), phosphorylated neurofilament (pNF) (Biolegend, SMI 31P, 1:500), Parvalbumin (Synaptic Systems, 195004 1:1000), GFAP (Sigma-Aldrich, G3893, 1:500), and OC (Sigma-Aldrich, AB2286, 1:5000). On the following day, slides were rinsed 3 times in PBS, incubated with fluorophore-conjugated secondary antibodies, and rinsed again 3 times in PBS. The following secondary antibodies were used: Donkey anti-rat IgG H+L Alexa Fluor 488 (Abcam, ab150153), Donkey anti-rabbit IgG H+L Alexa Fluor 647 (Abcam, ab150067), Donkey anti-mouse IgG H+L Alexa Fluor Plus 594 (Thermo Fisher Scientific, A32744) and Donkey anti-guinea pig IgG H+L Alexa Fluor 647 (Abcam, 706-605-148), all at 1:500 dilution. Finally, tissue sections were counterstained with DAPI (Thermo Fisher Scientific, A32733) and mounted onto glass slides using Fluoromount-G (Sigma-Aldrich, T5941). For thioflavin T staining, after secondary antibody staining, sections were incubated with 0.01% Thioflavin T (Sigma-Aldrich, T3516) in water for 5 minutes, followed by 50% ethanol and PBS washes. For the immunodetection of perineuronal nets (PNNs), after secondary antibody staining, slices were incubated overnight at 4 °C with a solution containing biotinylated *Wisteria floribunda* Lectin (WFA) (Vector Laboratories, B-1355-2, 1:300). On the following day, sections were rinsed 3 times in PBS (5 min each) at RT, incubated with a solution of red fluorescent streptavidin (Alexa Fluor™ 555 conjugate) (Thermo Fisher Scientifics, S32356, 1:300) and 5% NCS in PBS for 1h at RT, and rinsed again 3 times in PBS. Images were acquired using a Zeiss Axio Scan.Z1 slide scanner (20× objective). Quantification was performed using Arivis Vision4D software, employing standardized macros for image thresholding and segmentation. Regions of interest (ROIs) were manually drawn around the dentate gyrus, CA1, cortex, or the entire hippocampus. Quantification metrics included: total number of cFOS⁺ nuclei in the dentate gyrus; total number of WFA⁺ objects and the percentage of PV^+^ interneurons surrounded by PNNs in the CA1 region; number and total area of GFAP⁺ astrocytic objects; total pNF⁺ area in CA1; total number and area of IFITM3⁺ cells in cortex and hippocampus; amyloid plaque density, number, and average size of Thioflavin T⁺, 6E10⁺, and OC⁺ amyloid deposits.

### FACS sorting of microglia

Brains were collected and placed on ice in sterile phosphate-buffered saline (PBS). After removal of the meninges, cortical hemispheres were mechanically dissociated in Dounce buffer (15 mM HEPES, 0.5% glucose, HBSS 1x) using a tissue homogenizer (Fisher Scientific, 10198611). Myelin was removed by centrifugation through a 25% isotonic Percoll gradient at 800 x g for 15 minutes at 4°C. The resulting cell pellet was resuspended in PBS, centrifuged, and processed for flow cytometry to isolate microglia as previously described (Bhojwani-Cabrera et al., submitted manuscript). Briefly, prior to antibody staining, samples were blocked with Fc-block CD16/CD32 (Biolegend, 101320, 1:50) in FACS Buffer sterile filtered (1% FCS, 2mM EDTA, 25mM HEPES in PBS) for 10 min on ice and protected from light to block nonspecific binding. For cell surface staining, cells were resuspended in the appropriate antibody cocktail and incubated for 30 min on ice protected from light. Primary antibodies used were anti-mouse GFP (Aveslab, GFP-1020, 1:500), anti-mouse Cd11b (BioLegend, 101235, 1:500), anti-mouse CD45 (BioLegend, 103106, 1:500). Samples were centrifuged and washed with FACS buffer. Viability was assessed by staining with DAPI (1 µg/mL). Cells were sorted using a BD FACS Aria III cell sorter (BD Bioscience). To sort the cells, a 85-micron nozzle with 4-Way purity mode was used. The gating strategy involved selecting the cell population based on forward scatter area (FSC-A) to discriminate cells by size, and side scatter area (SSC-A) to separate them by granularity or internal complexity. Doublets were then excluded by gating FSC-height (FSC-H) and FSC-A to isolate single cells. All final analysis and data output were performed using FlowJo software (BD). For RNA-seq experiments, microglia were isolated from the whole adult mouse cortex, yielding 200,000-300,000 cells. Sorted cells were collected directly into RLT buffer (Quiagen, 79216) for subsequent RNA extraction.

### Real-time quantitative PCR (RT-qPCR)

To analyze gene expression in acutely FACS-isolated microglia from the adult mouse hippocampus, RNA was extracted using the RNeasy Mini Kit (Qiagen, 74104) with on-column DNase digestion (Thermo Fisher Scientific, 18047019). For BV2 cells, RNA was isolated using TRIzol reagent (Thermo Fisher Scientific, A33251). For all samples, RNA purity, integrity, and concentration were assessed by spectrophotometry (Nanodrop ND-1000 spectrophotometer) and micro-capillary electrophoresis (2100 Bioanalyzer, Agilent). Complementary DNA was synthesized using the RevertAid First Strand cDNA Synthesis Kit (Thermo Fisher Scientific, EP0442) following the manufacturer’s protocol, and quantitative PCR was performed using EvaGreen Master Mix (Cultek, 755032) on a QuantStudio 3 system (Applied Biosystems). Relative expression levels were analyzed using the ΔΔCt method (163), with Gapdh as the housekeeping gene for normalization. Primer sequences used in RT–qPCR assays are listed in Supplemental Table 6.

### Human transcriptomic samples (MSBB Cohort)

RNA-seq data from the parahippocampal gyrus (Brodmann area 36) were obtained from the Mount Sinai Brain Bank (MSBB) via the AMP-AD Knowledge Portal (Synapse ID: syn3159438; metadata: syn29855570; count data: syn27068754). We used processed raw count matrices in which genes with fewer than 1 count per million (CPM) had already been filtered out. For our analyses, we further subset the dataset to include only PHG samples of AD cases aged 60 to 89 years at death. Resequenced samples were handled by excluding unique sequencing batches; when biological replicates from the same individual were available, those with higher RNA integrity (RIN and RIN^2^ scores) were preferentially retained. To reduce sex-related confounding effects, samples were additionally filtered based on the expression of sex-linked genes, resulting in the removal of one sample (Supplemental Figure 1, A and B). The final dataset included 34 female and 32 male AD cases. For additional analyses, samples were filtered using the same criteria, selecting either control samples aged 60 to 89 years at death (19 females and 23 males). Statistical tests were carried out to assess data distribution prior analysis, we compared age at death and RIN across groups stratified by sex and disease status (Supplemental Figure 1, E and F). After observing significant differences in the distribution of age at death, we included age at death as a covariate in subsequent differential expression analyses.

### Mouse transcriptomic profiling by RNA-seq

For the Alzheimer’s disease-related samples (GSE304521), total RNA was extracted from hippocampi, rapidly dissected on ice using fine forceps. For the poly(I:C)-related samples (GSE304362), total RNA was extracted from acutely FACS-isolated microglia from the adult brain. For all samples, total RNA was extracted using the RNeasy Mini Kit (Qiagen, 74104), including on-column DNase digestion (Thermo Fisher Scientific, 18047019). RNA quality was assessed by micro-capillary electrophoresis (Agilent TapeStation, Agilent), and only samples with a RNA Integrity Number (RIN) ≥ 7 were used for library construction. Libraries were amplified by PCR, purified, and size-selected to enrich for fragments compatible with Illumina sequencing, followed by quality assessment using fluorometry (Qubit fluorometer, Thermo Fisher Scientific) and micro-capillary electrophoresis (Agilent TapeStation, Agilent). Libraries were prepared by Novogene Europe (Cambridge, UK) using a strand-specific protocol with poly-A selection and sequenced on an Illumina NovaSeq X Plus platform (PE150). For the Alzheimer’s disease-related samples (GSE304521), RNA-seq libraries were constructed using Novogene NGS Stranded RNA Library Prep Set (PT044). Raw paired-end RNA-seq reads in FASTQ format were processed by Novogene Europe (Cambridge, UK) using their standard bioinformatics pipeline. Briefly, adapter sequences, reads containing poly-N, and low-quality bases were removed using fastp (164). Quality metrics such as Q20, Q30, and GC content were also calculated during this step. All downstream analyses were based on the resulting high-quality clean reads. Clean reads were aligned to the mm39 mouse reference genome using HISAT2 (v2.2.1; (165)). The reference genome index was built using HISAT2, and gene model annotations were used to enable splice-aware alignment with improved accuracy. Library sizes of primary mapped reads were consistently above 45 million fragments. Gene-level quantification was performed using featureCounts (v2.0.6; (166)). The resulting count tables were used for downstream differential expression analysis. For the poly(I:C)-related samples (GSE304362), RNA-seq libraries were constructed using Novogene NGS RNA Library Prep Set (PT042). Raw paired-end data were processed in-house following a custom analysis pipeline. Briefly, raw paired-end RNA-seq reads in FASTQ format were first subjected to quality control using FastQC (v0.11.9; https://www.bioinformatics.babraham.ac.uk/projects/fastqc/) to assess base quality and sequence content. Adapter sequences and low-quality bases were then removed using Trim Galore (v0.6.7; https://zenodo.org/records/7598955) in paired-end mode. Post-trimming quality was re-evaluated with FastQC to confirm adapter removal and improved read quality. High-quality reads were aligned to the GRCm38 mouse reference genome using HISAT2 (v2.2.1; (167)) with the --dta option to facilitate downstream transcript assembly. Library sizes of primary mapped reads were consistently above 50 million fragments. Alignment output in SAM format was converted and sorted into BAM format using SAMtools (v1.13; (168)), and subsequently indexed. For genome browser visualization, BAM files were converted to TDF format using IGVTools (v2.5.3), aligned to the mm10 genome build, and visualized in IGV (169). Gene-level quantification was performed using HTSeq-count (v0.13.5) with parameters - -stranded=no, --type=exon, and --order=pos. Gene annotations were obtained from Ensembl release 102 (Mus_musculus.GRCm38.102.chr.gtf). The resulting count tables were used for downstream differential expression analysis.

### Differential gene expression analysis

Differential expression analyses were conducted in R version 4.5.0 (R Core Team, 2025) using DESeq2 (v1.48.1; (170)), considering genes with adjusted P value < 0.05 as significantly differentially expressed. Batch correction was applied when needed using the ComBat_seq() function from the sva package (v3.56.0, (171)). To facilitate the detection of batch effects, outliers, and major sources of variance, Principal Component Analyses were performed on variance-stabilized data (VST for datasets with >30 samples) or regularized log-transformed data (rlog for datasets with <30 samples) using the plotPCA() function from the DESeq2 package. The top 500 most variable genes were used for dimensionality reduction. The first two principal components were visualized with ggplot2 (v3.5.2, (172)), and sample labels were added using geom_text_repel() from ggrepel (v0.9.6, (173))to improve readability. Heatmaps were generated on gene expression counts rlog-transformed using pheatmap (v1.0.13, (174)), with hierarchical clustering applied to genes and samples ordered by experimental design. Rows were scaled using z-scores, and styling was harmonized with the overall plot design. Volcano plots were generated using ggplot2, where the –log₁₀ adjusted p-value was plotted against the log₂ fold change. Genes were color-coded by significance (padj < 0.05 or < 0.1) and classified as upregulated, downregulated, or not significant. Key interferon-alpha–related genes were labeled using ggrepel. Optional axis cutoffs were applied to reduce the impact of extreme outliers on plot scaling.

### Functional enrichment analysis

Functional enrichment analyses were performed on significantly differentially expressed genes as input (padj < 0.05) using Enrichr (175) and the MSigDB “hallmark” 2020 gene set collection (57). Gene Set Enrichment Analysis (GSEA) was performed to identify coordinated shifts in predefined gene sets, using the GSEA_4.4.0 desktop application (58), with the following parameters: 1,000 phenotype permutations, No_Collapse, and a fixed permutation seed (149). Ranked gene lists (.rnk files) were created from DESeq2 output using tidyverse tools (v2.0.0, (176))without additional filtering. Enrichment was tested against the MSigDB “hallmark” gene sets for both human (v2025.1.Hs) and mouse (v2025.1.Mm) (57) GSEA plots were recreated in R using ggplot2, cowplot (v1.2.0, (177)), and gridExtra (v2.3, (178)) to maintain stylistic consistency across visual outputs.

### Mouse behavioral testing

All behavioral assays were conducted during the light phase under consistent illumination (∼20 lux) in a dedicated testing suite. Mice were acclimated to the testing room for at least 30 minutes before each test. Behavior was recorded and analyzed using SMART 3.0 software (Panlab). We used adult mutant and control littermates of both sexes. For the open field (OF) test, mice were placed in the center of a 40 × 40 × 40 cm arena and allowed to explore freely for 15 minutes. Total distance traveled and time spent in the center zone were recorded as measures of locomotor activity and anxiety-like behavior, respectively. The *elevated plus maze* (EPM) consisted of two open and two closed arms (30 × 5 cm), elevated 50 cm above the floor. Mice were placed in the center facing an open arm and allowed to explore for 5 minutes. The percentage of time spent and number of entries into open arms were calculated. To evaluate spatial working memory spontaneous alternation was measured using the *Y-maze*. Briefly, the transparent symmetrical Y-maze consisted of three arms (30 × 5 × 15 cm). Three objects of similar size (∼20 cm3) were situated surrounding the maze at ∼15–20 cm outside the walls and mice were allowed to freely explore the maze for 8 minutes. An alternation was defined as sequential entry into all three arms (triad). Percent alternation = (Number of alternations / [Total arm entries − 2]) × 100. To study short-term spatial memory, we used the *T-maze*. The apparatus consisted of three arms, two of them situated at 180° from each other and the third situated at 90° with respect to the other ones representing the stem arm of the T. All three arms were 45 cm long. In the training trial, one arm was closed (novel arm) and mice were placed in the stem arm of the T (home arm) and allowed to explore this arm and the other available arm (familiar arm) for 10 min, after which they were returned to the home cage. After an inter-trial interval of 1 h mice were placed in the stem arm of the T-maze and allowed to freely explore all three arms for 5 min (testing phase). The time in the new arm∗100/total time in the two arms at 180° was the parameter evaluated. In the *novel object location* (NOLT) memory task, mice were habituated to a 40 × 40 cm arena for 15 minutes. Different distal visual cues were allocated in the walls of the arena. On day 2, two identical objects were presented during a 10-minute familiarization phase. After a 24-hour delay, one object was relocated to a novel position (test phase), and mice were allowed to explore for 5 minutes. The discrimination index was calculated as: (Time_novel − Time_familiar) / Total_exploration_time. Associative memory was assessed using *fear conditioning paradigm*. A 25 x 25 x 25 cm black chamber system was used (Panlab, Spain). On Day 1, animals were allowed to habituate for 5 minutes. After habituation, a tone (90 dB) was presented for 30 seconds. During the final 2 seconds of the tone, a 0.4 mA foot shock was administered for 1–2 seconds and the mouse was allowed to remain 30 additional seconds in the chamber. This procedure was repeated two times, and the mouse remained 2 additional minutes in the chamber before removing it. On Day 2, the contextual memory test was conducted. The mouse was placed in the chamber for 5 minutes, with no tone presented, and freezing behaviour was recorded. Cued memory was assessed on day 3 in a novel context, consisting of a 5-minute baseline followed by a 30-seconds tone presentation. Freezing was defined as the absence of all movement except respiration for >1 second. Animals were tracked and recorded with the Packwin V2.0.08 software (Panlab, Spain).

### Statistics

Data are presented as mean ± standard error of the mean (SEM), unless otherwise indicated. Statistical analyses were performed using GraphPad Prism (version 7) or R (version 4.2 or later). For comparisons between two groups, unpaired two-tailed Student’s t tests were used. Comparisons involving more than two groups were analyzed using one-way ANOVA followed by Bonferroni post hoc correction. Analyses involving two categorical factors used two-way ANOVA with Tukey’s HSD for multiple comparisons, and interaction terms were evaluated. For non-normally distributed data, the Mann– Whitney U test was applied. Correlations were assessed using Pearson’s correlation coefficient. A *P* value less than 0.05 was considered statistically significant. Specific statistical tests and sample sizes (n) for each experiment are reported in the figure legends.

### Study approval

The study was conducted in compliance with European Union Directive 2010/63/EU on the protection of animals used for scientific purposes. All the protocols for animal experimentation were approved by the Animal Welfare Committee at the Instituto de Neurociencias, the CSIC Ethical Committee, and the Dirección General de Agricultura, Ganadería y Pesca of Generalitat Valenciana. Mice were maintained under specific pathogen-free (SPF) conditions at the Animal Facility of the Instituto de Neurociencias (CSIC-UMH), on a 12 h light/12 h dark cycle (7:00 a.m. to 7:00 p.m.), at 20–24 °C and 40–60% controlled humidity, with *ad libitum* access to food and water.

### Data availability

The transcriptomics data sets generated in this study can be accessed at the GEO public repository using the accession number GSE304521 and GSE304362. In addition, human RNA-seq data used in this study were obtained from the publicly available The Mount Sinai Brain Bank (MSBB) study, hosted on the AD Knowledge Portal (https://adknowledgeportal.org/). The dataset is accessible via Synapse (https://www.synapse.org/): syn3159438; metadata: syn29855570; count data: syn27068754). The results published here are in whole or in part based on data obtained from the AD Knowledge Portal (https://adknowledgeportal.org/). These data were generated from postmortem brain tissue collected through the Mount Sinai VA Medical Center Brain Bank and were provided by Dr. Eric Schadt from Mount Sinai School of Medicine. Processed data and code used for statistical analysis, gene expression processing, and figure generation are available upon request from the corresponding author.

## Author contributions

Conceptualization, V.L-L., and J.P.L-A.; methodology, V.L-L., M.G-G., G.I., A.E-C., A.B-C., A.G., and J.P.L-A.; investigation, V.L-L., M.G-G., and A.E-C.; data curation, V.L-L., M.G-G., G.I., and M.G-F.; formal analysis: V.L-L., M.G-G., and G.I.; writing – original draft, V.L-L., and J.L-A.; writing – review & editing, V.L-L., G.I., M.G-G., A.E-C., A.B-C., M.G-F., A.G., and J.L-A.; funding acquisition, J.P.L-A.; resources, J.P.L-A.; supervision, J.P.L-A. The order of authorship was determined by overall contributions and approved by the authors for the final manuscript.

## Acknowledgments

We thank Eduardo Soriano, Alberto Pascual, Jose V. Sánchez-Mut, Berta L. Sánchez-Laorden, and Violeta Durán-Laforet for the critical reading of the manuscript. We are also grateful to members of the López-Atalaya laboratory for stimulating discussions throughout the course of this work. We thank the personnel of the Mouse Facility and the Microscopy and Omics Core Services at the Instituto de Neurociencias for their support and technical assistance. BioRender.com was used to create Figure 1, A and B, Figure 2A, Figure 3A, Figure 4A, Figure 5A, and Figure 6A. This study was supported by grant PID2021-129053OB-I00 funded by MCIU/AEI/10.13039/501100011033 and by ERDF, EU, and the Generalitat Valenciana, Conselleria d’Educació, Universitats i Ocupació (PROMETEO 2020/007) to J.P.L-A.; V.L-L. is the recipient of a 2021 neuroscience doctoral fellowship from the Fundación Tatiana Pérez de Guzmán el Bueno. G.I. holds a Formación de Personal Investigador (FPI) contract (CEX2021-001165-S-20-2), associated with “Severo Ochoa” grant CEX2021-001165-S from the MCIU/AEI/10.13039/501100011033. A.B-C. held a Formación de Personal Investigador (FPI) contract (PRE2019-090492), associated with grant RTI2018-102260-B-I00 awarded to J.P.L-A., and funded by MCIU/AEI/10.13039/501100011033 and by “ERDF A way of making Europe”. M.G-F. is the recipient of a Formación de Profesorado Universitario contract (FPU22/01375) funded by the Spanish Ministry of Science, Innovation and Universities (MICIU). The Institute of Neurosciences is a Severo Ochoa Center of Excellence (grant CEX2021-001165-S, funded by MCIU/AEI/10.13039/501100011033). This work was also funded by European Union NextGenerationEU/PRTR CNS2022 – 135391 (“Consolidación” program) and the Fundación Ramón Areces (CIVP21A7024) both to A.G. The project that gave rise to these results received the support of a fellowship from “la Caixa” Foundation (ID 100010434). The fellowship code is LCF/BQ/DR21/11880016 to M.G.-G.

## Supplemental Figures

**Supplemental Figure 1.**
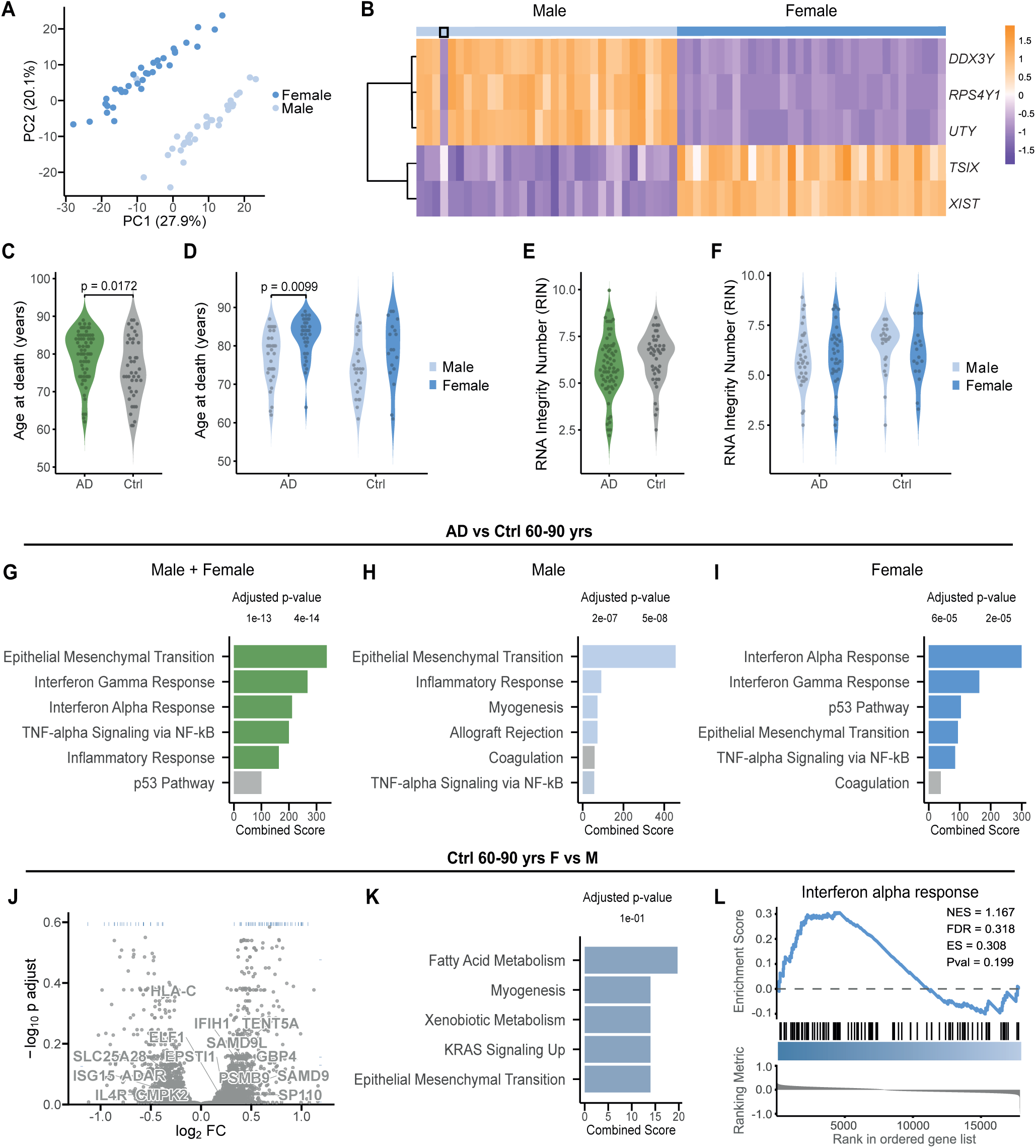
Increased interferon signaling in Alzheimer’s disease is sex-specific and absent in controls. (A) PCA of AD PHG RNA-seq data from the MSBB cohort (including samples from individuals aged 60–90 at death), with one misannotated male sample (excluded from analyses) clustering with female samples. (B) Heatmap of canonical sex chromosome gene expression in PHG from AD patients aged 60 to 90 years, confirming accurate sex annotation except for the misannotated male sample (bold). n = 32 (male AD), 34 (female AD). (C-F) Violin plots of age at death (C–D) and RNA integrity number (RIN; E–F) in AD and control samples, with stratification by sex where indicated. Two-group contrasts (C, E) were tested with the Mann–Whitney U test (Wilcoxon rank-sum) with continuity correction; sex × condition effects (D, F) were tested with two-way ANOVA followed by Tukey’s HSD. Results: age differed between AD and controls (C; P = 0.0172) and showed a sex difference within AD (D; Tukey’s HSD, P = 0.0099). RIN showed no significant differences either between AD and controls (E) or by condition, sex, or their interaction (F). (G–I) Functional enrichment of differentially expressed genes (FDR < 0.05) in PHG tissue of AD patients (60 to 90 years). (G) Upregulated pathways in AD versus age-matched control samples. (H) Upregulated pathways in male AD versus age-matched male controls. (I) Upregulated pathways in female AD versus age-matched female controls. Female AD samples exhibit stronger interferon signatures, whereas in male AD samples the enrichment of epithelial– mesenchymal transition and inflammatory response pathways are prominent. (J) Volcano plot of differential gene expression in PHG of control individuals (female vs. male, aged from 60 to 90 years), showing no significant changes in interferon-related genes in females (gray, non-significant). n = 23 (male AD), 19 (female AD). (K) Functional enrichment analysis of differentially expressed genes (FDR < 0.05) in the PHG from control individuals (female vs. male, aged from 60 to 90 years), showing no significant activation of interferon alpha or gamma response pathways. n = 23 (male AD), 19 (female AD). (L) GSEA plot of the interferon alpha response pathway in control individuals (female vs. male, aged from 60 to 90 years). n = 23 (male AD), 19 (female AD).

**Supplemental Figure 2.**
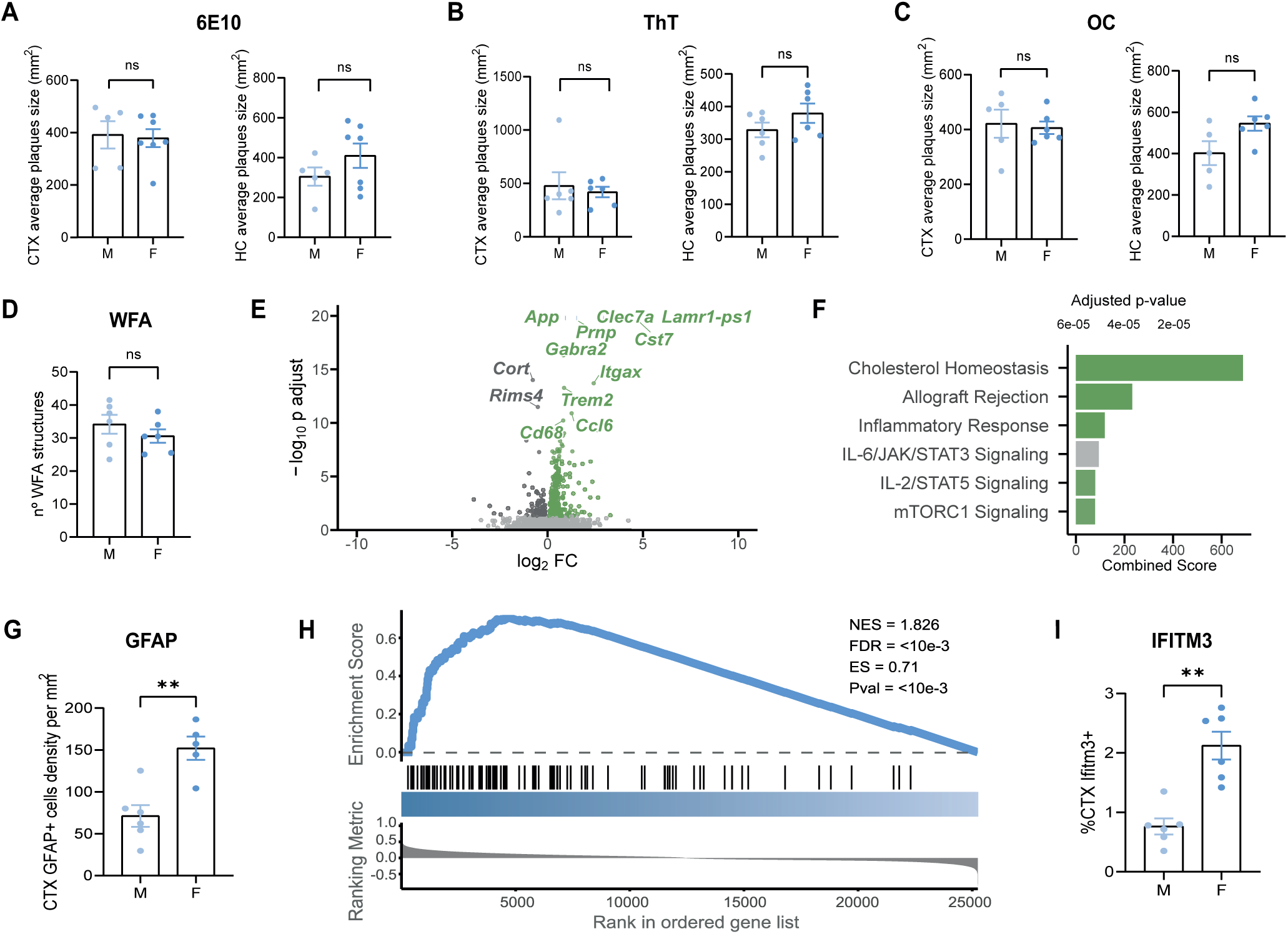
Sex-dependent neuropathological and transcriptomic changes in APP/PS1 Mice. (A) Quantification of Aβ plaque size by 6E10 immunohistochemistry in the cortex (CTX) and hippocampus (HC) of 6-month-old male and female APP/PS1 mice, showing no significant differences in plaque size between sexes (ns, *P* > 0.05, Mann-Whitney test). (B) Quantification of fibrillar Aβ deposit size by Thioflavin T staining in CTX and HC, confirming no significant differences in deposit size between sexes (ns, *P* > 0.05, Mann-Whitney test). (C) Quantification of β-sheet-rich Aβ conformation size by OC antibody staining in CTX and HC, showing no significant differences between sexes (ns, *P* > 0.05, Mann-Whitney test). (D) Quantification of WFA-positive structure number in CA1, showing no significant differences between (ns, *P* > 0.05, Mann-Whitney test). (E) Volcano plot of differential gene expression in HC contrasting APP/PS1 vs control, highlighting top 15 significant genes (green, *P* < 0.05 in APP/PS1; dark grey, *P* < 0.05 in control; gray, non-significant). (F) Functional enrichment analysis from HC tissue of mice showing enriched pathways for APP/PS1 mice contrasted against control mice, where no interferon pathway was highlighted. (G) Quantification of GFAP immunohistochemistry density in the cortex, showing increased astrogliosis in female APP/PS1 mice (*\*\*P* ≤ 0.01), Mann-Whitney test). (H) GSEA plot of the interferon alpha response pathway in the hippocampi of female vs. male APP/PS1 mice. (I) Quantification of IFITM3 immunohistochemistry intensity in the cortex, showing increased labeling in female APP/PS1 (*\*\*P* ≤ 0.01), Mann-Whitney test). Data from 6-month-old APP/PS1 mice. IHC: n = 7 (control), 4 (Poly(I:C) 24h), 4 (Poly(I:C) 72h), and 6 (APP/PS1). IHC: n = 12 (6 females, 6 males). RNA-seq: n = 8 AD females, 8 AD males, 8 WT females, 8 WT males. Data represent mean ± SEM.

**Supplemental Figure 3.**
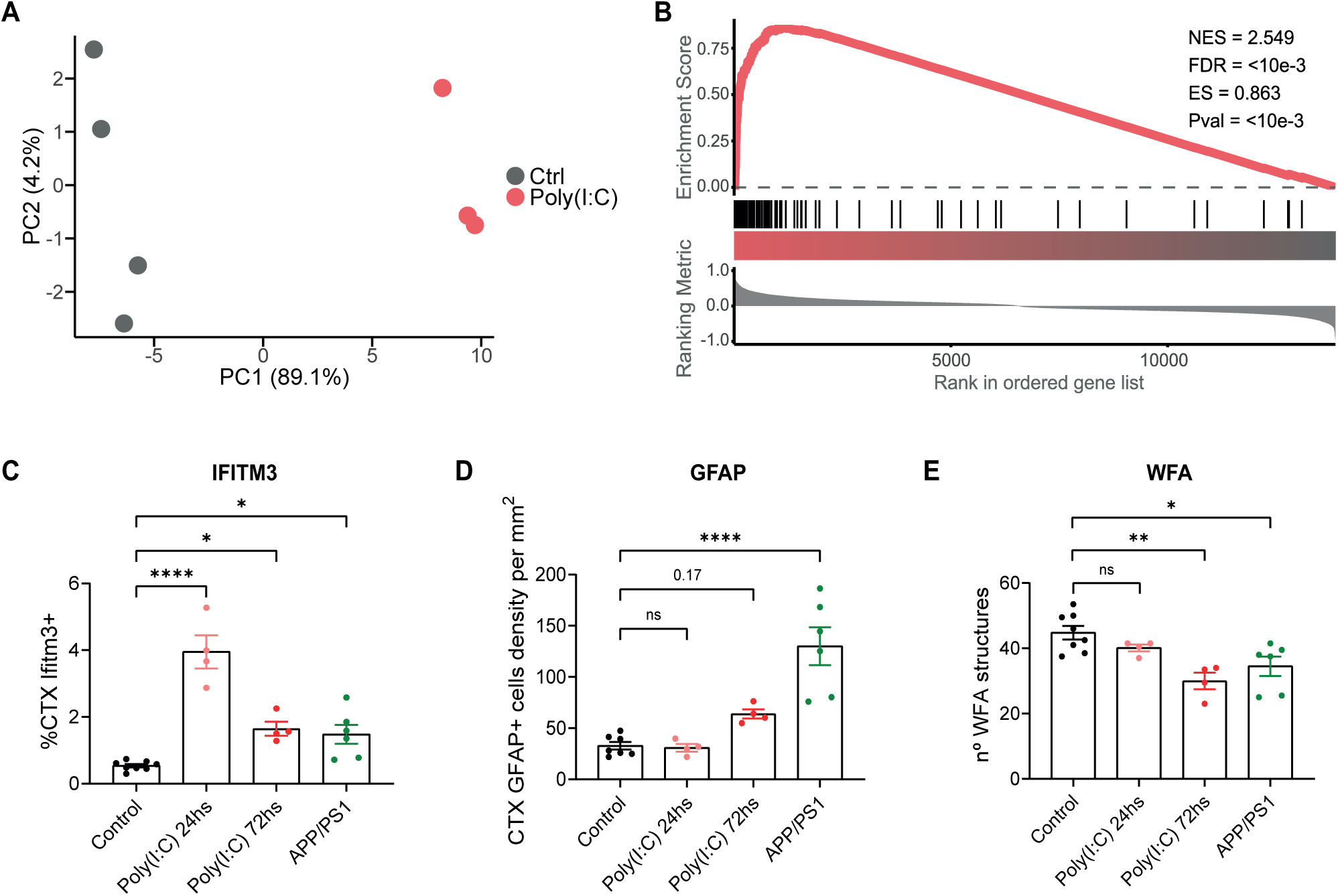
Poly(I:C)-induced interferon signaling and neuropathological changes in control mice. (A) PCA of hippocampal RNA-seq data from 3-month-old wild-type C57BL/6 mice treated with poly(I:C) (12 mg/kg) and analyzed at 24 hours, compared to untreated controls, showing transcriptomic divergence among the two groups. (B) GSEA plot of the interferon alpha response pathway in the HC of poly(I:C)-treated mice (24 hours) vs. untreated controls, confirming significant enrichment of interferon alpha signaling. (C) Quantification of IFITM3 immunohistochemistry intensity in the cortex, showing increased labeling in poly(I:C)- treated mice at 24 hours (*\*\*\*\*P* < 0.0001), and 72 hours (*\*P* < 0.05), and in APP/PS1 mice (*\*P* < 0.05) compared to controls (Mann-Whitney test). (D) Quantification of GFAP immunohistochemistry intensity in the cortex, indicating no significant changes in astrogliosis in poly(I:C)-treated mice at 24 hours (ns, *P* > 0.05) and 72 hours (ns, *P* > 0.05), and an increased in APP/PS1 mice (\*\*\*\**P* < 0.0001) compared to controls (Mann-Whitney test). (E) Quantification of WFA-positive structure number in CA1, showing no significant changes in poly(I:C)-treated mice at 24 hours (ns, *P* > 0.05), and fewer structures at 72 hours (*\*\*P* < 0.01) and APP/PS1 mice (*\*P* < 0.05) compared to controls (Mann-Whitney test). Data from 3-month-old wild-type C57BL/6 mice (RNA-seq at 24 hours, immunohistochemistry at 24 and 72 hours) and 6-month-old APP/PS1 mice. IHC: n = 7 (control), 4 (Poly(I:C) 24h), 4 (Poly(I:C) 72h), and 6 (APP/PS1). Poly(I:C) groups included only males; other groups were sex-balanced (3 females, 3 males for APP/PS1; 3 females, 4 males for controls). RNA-seq: n = 4 (control), 3 (Poly(I:C) 24h). Data represent mean ± SEM.

**Supplemental Figure 4.**
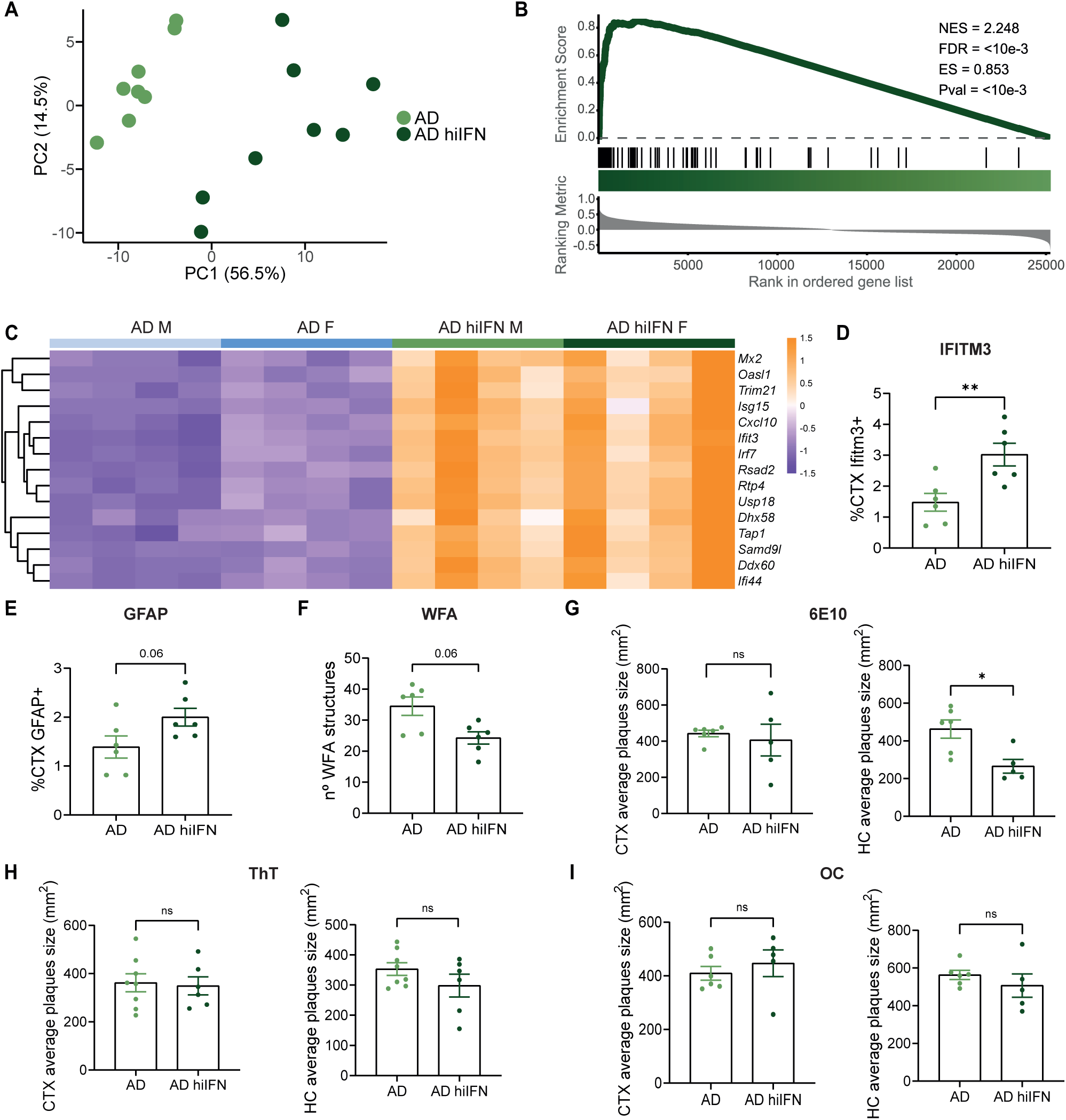
Neuropathological and transcriptomic effects of enhanced interferon signaling in APP/PS1 mice. (A) PCA of hippocampal RNA-seq data from 6-month-old APP/PS1 (AD) and microglia-specific *Rela* knockout (AD hiIFN) mice, showing transcriptomic divergence between the two groups. (B) GSEA plot of the interferon alpha response pathway in the hippocampus of AD hiIFN vs. AD mice, confirming significant enrichment of interferon alpha signaling. (C) Heatmap of interferon-alpha-related gene expression in hippocampal tissue from male and female AD and AD hiIFN mice, not highlighting a sex-biased transcriptomic profile. (D) Quantification of IFITM3 immunohistochemistry intensity in the cortex, showing higher labeling in AD hiIFN vs. AD mice (\*\**P* < 0.01, Mann-Whitney test). (E) Quantification of GFAP immunohistochemistry intensity in the cortex, indicating nearly significant increase in astrogliosis in AD hiIFN vs. AD mice (*P*= 0.06, Mann-Whitney test). (F) Quantification of 6E10 immunohistochemistry for Aβ plaque size in the cortex (CTX) and hippocampus (HC), showing reduced plaque size in AD hiIFN vs. AD mice hippocampus (\**P* < 0.05) and no significant changes in cortex (ns, *P* > 0.05) (Mann-Whitney test). (G) Quantification of Thioflavin T staining for fibrillar Aβ deposit size in CTX (ns, *P* > 0.05) and HC (ns, *P* > 0.05), showing no significant changes in AD hiIFN vs. AD mice (Mann-Whitney test). (H) Quantification of OC staining for β-sheet-rich Aβ conformation size in CTX (ns, *P* > 0.05) and HC (ns, *P* > 0.05), showing no significant changes in AD hiIFN vs. AD mice (Mann-Whitney test). Data from 6-month-old AD hiIFN and APP/PS1 mice. IHC: n = 6 per group (3 females, 3 males). RNA-seq: n = 8 per group (4 females, 4 males). Data represent mean ± SEM.

**Supplemental Figure 5.**
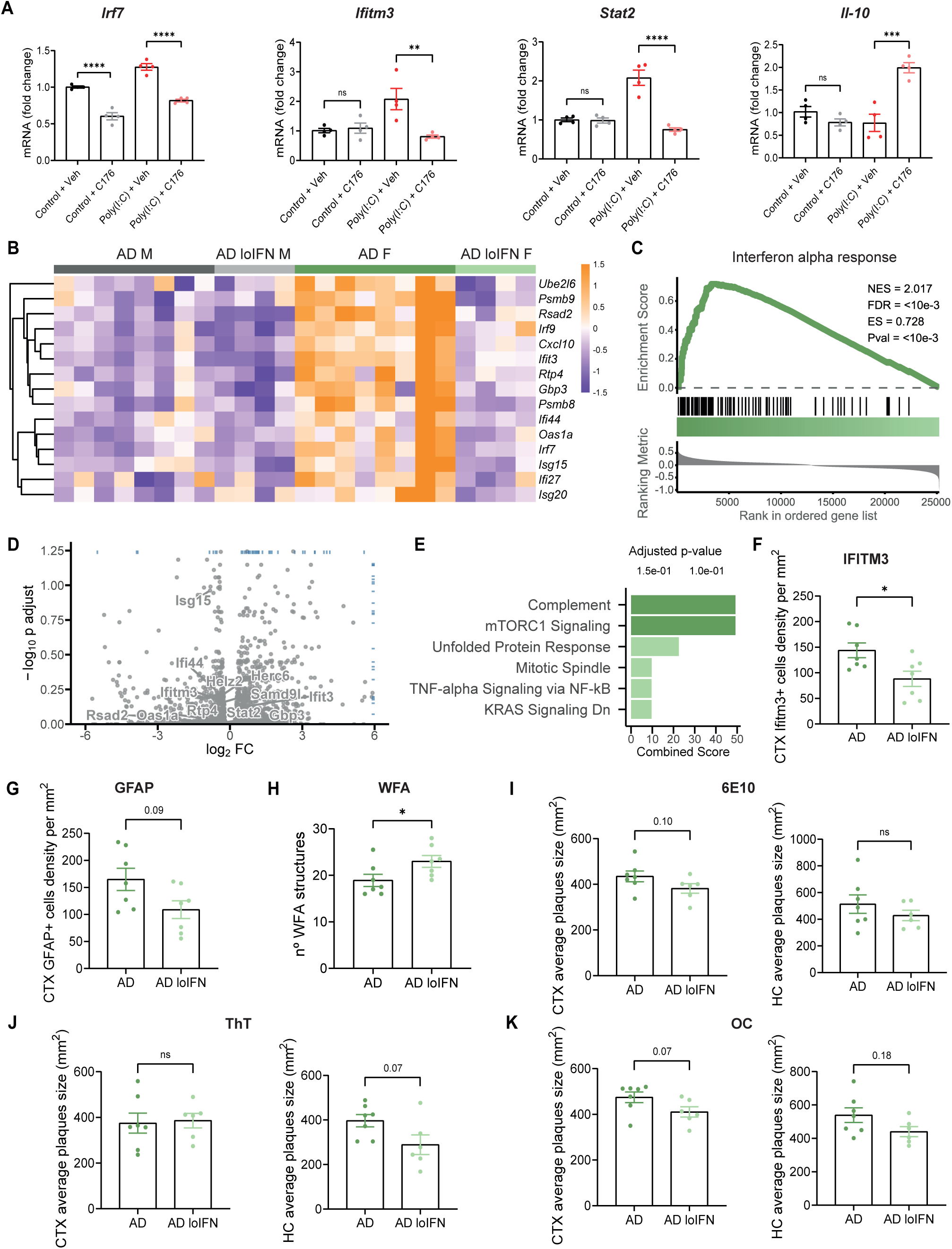
Effects of STING inhibition on BV2 microglial cells and neuropathology in APP/PS1 mice. (A) Expression of interferon-stimulated genes (*Irf7*, *Ifitm3*, *Stat2*) and the anti-inflammatory cytokine *Il10* in BV2 microglial cells treated with vehicle (Veh) or STING inhibitor C-176, with or without poly(I:C) stimulation (ns, *P* > 0.05, \**\*P* < 0.01, \*\*\**P* < 0.001, \*\*\**\*P* < 0.0001, Mann-Whitney test). (B) Heatmap of interferon-alpha-related gene expression in hippocampal tissue from male and female 6-month-old APP/PS1 (AD) and C-176-treated APP/PS1 (AD loIFN) mice, highlighting a higher transcriptomic profile switch in females. (C) GSEA plot of the interferon alpha response pathway from HC tissue of treated vs. untreated female AD mice, confirming reduced enrichment of interferon signaling in treated females. (D) Volcano plot of differential gene expression in the hippocampus of male AD loIFN vs. untreated AD mice, showing no significant changes in interferon-related genes (gray, non-significant). (E) Functional enrichment analysis from HC tissueof male AD loIFN vs. untreated AD mice, showing no significant reduction of the interferon alpha or gamma response pathways. (F) Quantification of IFITM3 immunohistochemistry intensity in the cortex, showing lower labeling in untreated AD mice vs. AD loIFN (\**P* < 0.05, Mann-Whitney test). (G) Quantification of GFAP immunohistochemistry intensity in the cortex, indicating nearly significance decrease of astrogliosis in AD loIFN vs. untreated AD mice (*P* = 0,09, Mann-Whitney test). (H) Quantification of WFA-positive structure number in CA1, showing an increase in AD loIFN vs. untreated AD mice (\**P* > 0.05, Mann-Whitney test). (I) Quantification of 6E10 immunohistochemistry for Aβ plaque size in the cortex (CTX) and hippocampus (HC), showing no significant changes in AD loIFN vs. untreated AD mice (ns, *P* > 0.05, Mann-Whitney test). (J) Quantification of Thioflavin T staining for fibrillar Aβ deposit size in CTX and HC, showing no significant changes in AD loIFN vs. untreated AD mice (ns, *P* > 0.05, Mann-Whitney test). (K) Quantification of OC staining for β-sheet-rich Aβ conformation size in CTX (*P* > 0.05) and HC (ns, *P* > 0.05), showing no significant changes in plaques size in AD loIFN vs. untreated AD mice (Mann-Whitney test). Data from BV2 cell cultures and 6-month-old AD loIFN and APP/PS1 mice. qPCR: n = 4 per group. IHC: n = 6 per group (3 females, 3 males). RNA-seq: n = 8 AD (females), 4 AD loIFN (females). Data represent mean ± SEM.

**Supplemental Figure 6.**
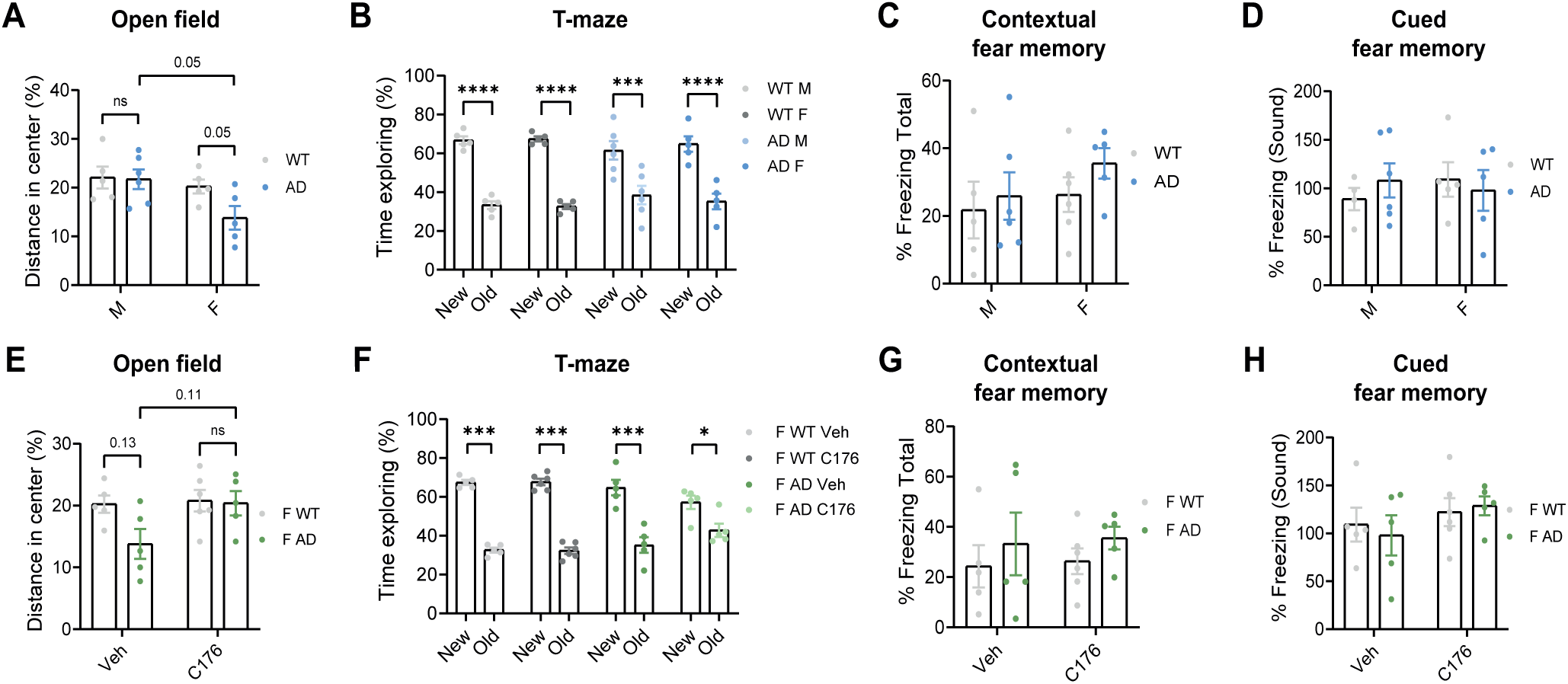
Behavioral tasks with no significant differences by sex or STING inhibition in APP/PS1 mice. (A) Percentage of distance traveled in the center zone of the open field test in 6-month-old male and female WT and APP/PS1 (AD) mice, showing reduced center exploration in AD females compared to males and WT controls (*P* = 0.05, two-way ANOVA with Tukey’s post hoc test). (B) Percentage of time spent exploring in the T-maze test, showing no significant differences between sexes or genotypes (\*\*\**P* < 0.001, \*\*\**\*P* < 0.0001, two-way ANOVA with Tukey’s post hoc test.) (C) Percentage of freezing during the contextual phase of fear conditioning, showing no significant differences across groups (ns, *P* > 0.05, two-way ANOVA with Tukey’s post hoc test). (D) Percentage of freezing during the auditory cue in cued fear conditioning, also showing no significant differences (ns, *P* > 0.05, two-way ANOVA with Tukey’s post hoc test). (E) Percentage of distance traveled in the center zone of the open field test in female AD mice treated with vehicle or C-176, showing a non-significant trend toward increased center exploration in C-176–treated mice (P = 0.11, two-way ANOVA with Tukey’s post hoc test). (F) Percentage of time spent exploring in the T-maze test in female AD mice, showing no difference between vehicle- and C-176– treated groups (ns, *P* > 0.05, two-way ANOVA with Tukey’s post hoc test). (G) Percentage of freezing during contextual fear conditioning in female AD mice, showing no effect of C-176 treatment (ns, *P* > 0.05, two-way ANOVA with Tukey’s post hoc test). (H) Percentage of freezing during cued fear conditioning in female AD mice, showing no effect of C-176 treatment (ns, *P* > 0.05, two-way ANOVA with Tukey’s post hoc test). All behavioral tests were performed in 6-month-old APP/PS1 mice. n = 5 (male WT), 6 (male AD), 5 (female WT), 5 (female AD), 5 (female WT + vehicle), 5 (female WT + C- 176), 5 (female AD + vehicle), and 5 (female AD + C-176). Data represent mean ± SEM.

## Notes

Conflict of interest: The authors have declared that no conflict of interest exists.

### Competing Interest Statement

The authors have declared no competing interest.

### Summary of Updates

Updated analyses related to Figure 1 and Supplemental Figure 1. The corresponding text has been updated. The main text has been refined for clarity and style.

